# Multi-scale gene regulatory networks for resolving transcription factor and autoimmune variant impacts on human B cell fate dynamics

**DOI:** 10.1101/2025.04.23.649973

**Authors:** Nicholas A Pease, Jingyu Fan, Swapnil Keshari, Jered Stratton, Steven B Gierlack, Akanksha Sachan, Zarifeh Heidari Rarani, Tanush Swaminathan, Christopher S McGinnis, Wenxi Zhang, Deepa Bisht, Trirupa Chakraborty, Peter Gerges, Godhev Manakkat Vijay, Louis Lau, Camila Macedo, Diana Metes, Hyejung Won, Ansuman T Satpathy, Abhinav K Jain, Nidhi Sahni, Jishnu Das, Harinder Singh

## Abstract

Bifurcation of activated human B cells into plasmablast (PB) and germinal center precursor (preGC) fates underlies protective and autoimmune antibody responses, yet gene regulatory networks (GRNs) governing the alternative trajectories remain poorly defined. Using temporal single-cell multiomics, we assembled state-specific human B cell GRNs spanning four scales: transcription factor (TF)-to-fate, TF-to-gene, cis-regulatory element (CRE)-to-gene and nucleotide-to-CRE. Applying the framework to *in vitro* differentiated and tonsil B cells revealed concordant regulatory architectures and impacts of *in silico* TF perturbations. CRISPR perturbations validated many TF-to-fate and TF-to-gene predictions and revealed a mutually repressive BATF-IRF4/PRDM1 network module. The GRNs were used to predict and interpret effects of autoimmune disease variants, uncovering partitioning of disease risk at distinct TF motifs and B cell states. Predicted functional variants were independently supported by chromatin and expression QTLs and reporter assays. A web application enables exploration of B cell regulatory determinants of vaccine responses and autoimmune diseases.

## Introduction

Bifurcation of activated B cells into plasmablast (PB) and germinal center precursor (preGC) trajectories is a critical determinant of antibody response quality, underlying both protective immunity and autoimmune pathogenesis^1–3^. PBs are short-lived cells that secrete lower-affinity antibodies, whereas preGC B cells delay terminal differentiation and initiate germinal center reactions where their progeny undergo somatic hypermutation and affinity-based selection through interactions with T follicular helper cells, ultimately generating memory B cells and long-lived plasma cells that encode higher-affinity antibodies^4^. The quality and durability of antibody responses depend on the balance between the generation of PBs and preGCs from activated B cells.

We and others have elucidated a small-scale gene regulatory network (GRN), comprising sequential reciprocal negative feedback loops coupled with an incoherent feed-forward loop between the transcription factors (TFs) IRF4-IRF8 and BLIMP1-BCL6 that control the generation of murine PB and preGC states^5–8^. However, this network represents only a subset of the TFs and regulatory linkages that constitute the much larger GRNs underlying B cell activation and fate bifurcation. Furthermore, previous B cell GRNs lack (i) temporal resolution across naïve, activated and differentiated cell states, (ii) delineation of cis-regulatory element (CRE)-to-gene linkages and (iii) base-pair resolution modeling needed to infer TF binding and activity at specific genomic sequences. Consequently, these GRNs are inherently incapable of predicting or interpreting the impact of non-coding genomic variants, which constitute 90% of autoimmune disease GWAS associations. To address these limitations and comprehensively analyze the impact of TFs, CREs and autoimmune variants on human B cell fate dynamics, we assembled temporally resolved human B cell GRNs at multiple scales linking TFs to fates, TFs to genes and CREs, CREs to genes, and nucleotides to CREs.

Single-cell transcriptional profiling coupled with perturbations and clonal tracking demonstrated that the human B cell fate bifurcation is manifested through a BCL6-dependent preGC state and an IRF4/BLIMP1-dependent PB state. By integrating temporally resolved single-cell multiomics with computational modeling and CRISPR perturbations, we executed a ‘Map-Predict-Perturb’ workflow to rigorously assemble and test B cell state-specific GRNs. We leveraged these multi-scale GRNs to predict and interpret the effects of thousands of autoimmune disease variants on TF activity, CRE accessibility and target gene expression. We provide a UCSC genome browser session (https://tinyurl.com/BcellMultiomeTracks) and a programming-free, interactive web application (https://pitt-csi.shinyapps.io/humanBcellGRN/) to explore the human B cell GRNs and their accompanying multi-scale predictions. This resource will facilitate comprehensive analysis of TFs and non-coding variants that control or impact B cell fate dynamics in the context of infection, vaccination and autoimmunity.

## Results

### Human B cells activated *in vitro* bifurcate into PB and preGC trajectories

While plasmablast (PB) differentiation has been extensively studied *in vitro*, the alternative germinal center (GC) B cell trajectory has received less attention^9–11^. To enable analysis of GRNs and molecular mechanisms that govern B cell fate choice, we adapted an existing culture system involving activation and differentiation of naïve human B cells to enhance the generation of both PB and GC precursor (preGC) trajectories^9^. Naïve B cells (>95% purity, **Extended Data Fig. 1a**) were activated for 4 days with anti-IgM, CpG, CD40L, and IL-2, then cultured for 2 days with CD40L, IL-2, IL-4, and IL-10 to enable the generation of both B cell trajectories (**Fig. 1a**). We tested for differentiation into the divergent trajectories by analyzing the reciprocal expression of IRF8 and IRF4, which are transcription factors (TFs) that demarcate preGC versus PB fates in murine B cells^5^ and show similarly divergent expression in human B cells responding to immunization with a flu vaccine^1^. At two days post-activation *in vitro*, most B cells uniformly induced expression of IRF4 and up-regulated IRF8 (**Fig. 1a**). By Day 4, most cells had undergone 1-4 cell divisions, and IRF4/IRF8 levels began to diverge. By Day 6, cells were bifurcated into distinct IRF8^lo^IRF4^hi^ and IRF8^hi^IRF4^lo^ states. To further validate PB and preGC emergence, we analyzed expression of BCL6, BLIMP1 and AID, which are also reciprocally expressed in preGC and PB states^5^. The IRF8^lo^IRF4^hi^ cells were BCL6^lo^BLIMP1^hi^ and AID^lo^, whereas IRF8^hi^IRF4^lo^ cells were BCL6^hi^BLIMP1^lo^ and AID^hi^, consistent with these two populations representing PB and preGC trajectories, respectively (**Fig. 1b,c**). In keeping with these results, increasing CD40L, which mimics sustained T cell help, enhanced preGC versus PB differentiation (**Extended Data Fig. 1b**). Notably, the generation of both trajectories was highly reproducible across 8 different donors (4 female and 4 male) representing diverse ethnicities and ages (**Extended Data Fig. 1c**). Isotype switching occurred at similar frequencies in both populations (9% vs. 5%, **Extended Data Fig. 1d**), ruling out class switching as a major fate modulator.

**Fig. 1.**
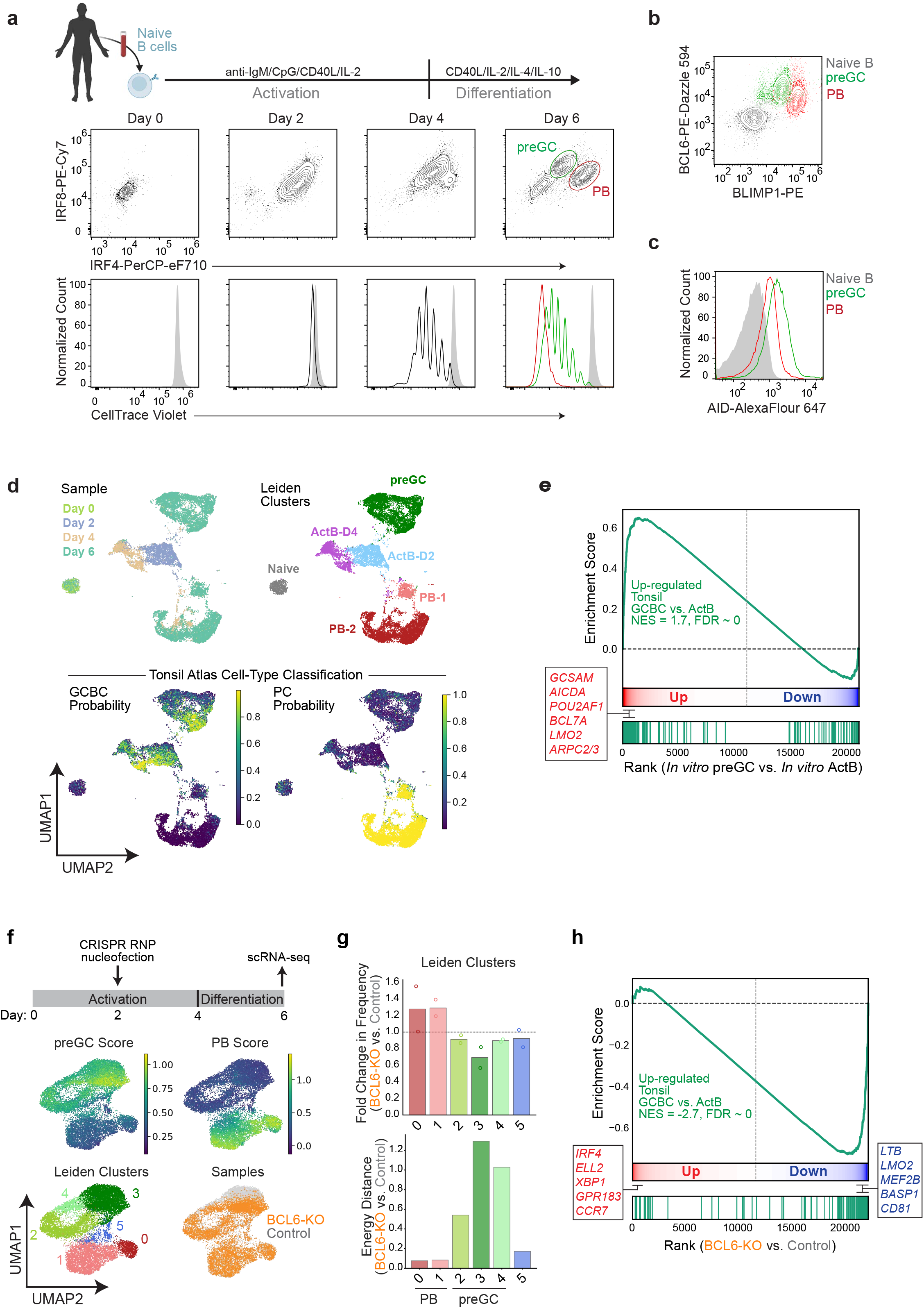
Human B cells activated *in vitro* bifurcate into PB and preGC trajectories. **(a)** Naïve B cells, isolated from healthy human donor PBMCs, were labeled with CellTrace proliferation dye and stimulated *in vitro* as indicated in the schematic (*top*) (see Methods). IRF4 and IRF8 expression dynamics were monitored by intracellular flow cytometry at indicated days (*middle*). Cell proliferation was monitored by CellTrace dilution at each timepoint (*bottom*). **(b)** Flow cytometry analysis of BLIMP1 and BCL6 expression in preGC (green) and PB (red) populations at Day 6. Naïve B cells (gray) from Day 0 are overlaid for reference. **(c)** Flow cytometry analysis of AID expression in indicated populations as in panel (b). **(d)** Single-cell RNA-seq profiling of *in vitro* activated B cells at D0, D2, D4, and D6. Cells in the aggregated UMAP are colored by timepoints (*top left*), Leiden clusters (*top right*), and support vector machine classification probabilities based on Human Tonsil Atlas B cell states (*bottom*). **(e)** Gene set enrichment analysis using top 150 most up-regulated genes (vertical green lines) between tonsil GCBC and activated B cell (ActB) populations. Genes were ranked according to their differential expression score between *in vitro* preGC and ActB populations from Fig. 1d. Inset highlights select tonsil GCBC-associated genes. **(f)** Naïve B cells were activated for 2 days as in Fig. 1a before nucleofection with CRISPR RNP complexes containing either non-targeting control gRNAs (control) or BCL6 gRNAs (BCL6-KO) (n = 2 donors) at D2. Cells were re-cultured according to differentiation protocol above and harvested at D6 for sample hash-tagging and scRNA-seq. Control and BCL6-KO cells from both donors were co-embedded in an aggregated UMAP displaying preGC and PB gene scores (see Methods), Leiden clusters, and sample (BCL6-KO vs. Control). **(g)** State-specific comparison of BCL6-KO vs. Control cell frequencies and transcriptional states. Bar plot displays fold change in cluster frequency between BCL6-KO and Control cells (n = 2 donors) (*top*). Bar plot displays energy distance between BCL6-KO and control cells within each Leiden cluster (*bottom*). **(h)** Gene set enrichment analysis using top 150 most up-regulated genes (vertical green lines) between tonsil GCBC and ActB populations. Genes were ranked by differential expression score between BCL6-KO and control cells in the preGC Leiden cluster (cluster 3). Inset highlights select tonsil GC-associated genes that are BCL6- dependent.

To comprehensively analyze the distinct B cell trajectories and examine the correspondence of their transcriptional programming with *in vivo* PB and GCBC states, we performed single-cell RNA-seq (scRNA-seq) at Days 0, 2, 4, and 6. We used probabilistic topic modeling to represent each cell as a weighted mixture of gene expression programs, reducing feature complexity while preserving biological interpretability^12^. Low-dimensional UMAP representations revealed that Day 2 cells represented a relatively homogeneous transcriptional state that was well resolved from naïve B cells (**Fig. 1d**). By Day 4, cells began to manifest divergent transcriptional states that became fully populated by Day 6, mirroring the bifurcated states observed by expression of reciprocally regulated TFs (**Fig. 1a** and **Extended Data Fig. 1e**, see **Supplementary Table 1** for complete marker gene analysis). To objectively classify the *in vitro* cell states, we trained a support vector machine (SVM)^13^ model on tonsillar B cell populations from the Human Tonsil Atlas^14^ (**Fig. 1d**). These classification results showed a bifurcated pattern of GCBC or plasmablast/plasma cell (PB/PC) probabilities among Day 6 cells, supporting the correspondence of the *in vitro* B cell states with their *in vivo* counterparts.

To confirm that the Day 6 preGC population adopts a transcriptional program that is distinct from activated B cell states from which it is derived, we performed gene set enrichment analysis with a gene set that distinguishes the tonsil GCBC transcriptome from that of tonsil activated B cells. The top 150 most up-regulated genes in tonsil GCBCs (compared with tonsil activated B cells) were highly enriched among genes that were most up-regulated in our *in vitro* preGC cluster (compared with *in vitro* activated B cell clusters) (**Fig. 1e**). Many of these genes have known roles in marking and/or regulating GCBC functionality (e.g. *GCSAM*, *LMO2*, *BCL7A*, *AICDA*, *APRC2/3*, *POU2AF1*). These results, along with differential expression of additional marker genes (**Supplementary Table 1**), demonstrate the generation of divergent PB and preGC states that are distinguishable from activated B cells in the *in vitro* system.

Paired B cell receptor sequencing (BCR-seq) of Day 6 cells revealed that individual naïve B cell clones can give rise to both PB and preGC trajectories (n=16) (**Extended Data Fig. 1f**), demonstrating that B cell fate choice can occur after the first cell division as previously shown for murine B cells *in vivo*^15^. However, the observed number of PB-only and preGC-only clones significantly exceeded the number of expected single-fated clones, under the assumption that clonal sibling fates are completely independent (n=641 PB-only (p < 2e-16) and n=38 GC-only (p < 2e-16)). These data support a model in which, following fate specification, differentiated states are stably propagated in clonal progeny, as observed for murine PB differentiation^16,17^. In agreement with this model, the observed number of bifurcated clones was lower than expected (n=16, p = 5e-12). Nonetheless, the presence of bifurcated clones suggested that fates of individual B cell clones are not pre-determined but emerge independently across and within clonal families. In conjunction with the initial manifestation of divergent transcriptional states at D4 (**Fig. 1d**), these results suggest that activated B cells probabilistically diverge into PB or preGC states which are then heritably propagated.

Since current *in vitro* GC systems cannot support bona fide GCBC formation and dynamics, we asked instead whether the *in vitro* preGC trajectory shares the transcriptional\ requirement for BCL6 that underlies GCBC initiation *in vivo*^18,19^. To test the requirement of BCL6 in preGC transcriptional programming, we used CRISPR/Cas9 editing to knockout (KO) BCL6 at Day 2 and performed scRNA-seq analyses of the control and perturbed cells at Day 6 (**Fig. 1f**). CRISPR-induced indel generation at the BCL6 locus and reduced BCL6 protein levels were confirmed by PCR genotyping and flow cytometry, respectively (**Extended Data Fig. 1g,h**). As expected, the integrated analysis of control and BCL6-KO cells revealed two major populations corresponding to preGC and PB trajectories (**Fig. 1f**), consistent with the bifurcation observed by flow cytometry (**Fig. 1a**). While both control and BCL6-KO cultures contained preGC and PB populations (**Fig. 1f**), BCL6-KO cells generated more PBs and fewer preGCs, consistent with the known role of BCL6 in antagonizing PB formation in favor of the GC trajectory (**Fig. 1g**). Although BCL6-KO caused only moderate decreases in the frequency of cells in preGC clusters, BCL6 deficiency profoundly disrupted the preGC transcriptional state when analyzed at higher resolution (**Fig. 1g**). For each cluster, we measured the difference between BCL6-deficient cells and control cells in a high-dimensional manner by calculating the energy distance –a metric summarizing between-condition versus within-condition distances in gene expression space^20^. The energy distance was highest in preGC clusters, demonstrating that BCL6 loss specifically disrupts the preGC transcriptional state without affecting PB differentiation as expected (**Fig. 1g**). To investigate whether BCL6-deficient preGC cells specifically failed to establish a GCBC transcriptional program, we performed gene set enrichment analysis using the tonsil GCBC versus tonsil activated B cell gene set. The GCBC-specific up-regulated genes were significantly enriched among those down-regulated in BCL6-KO versus control preGC cells (NES = −2.7, FDR ∼0), including genes known to mark and/or promote GCBC formation such as *BASP1*, *CD81*, *LTB*, *LMO2*, and *MEF2B*^21–25^ (**Fig. 1h**). Conversely, BCL6-deficient cells expressed higher levels of transcriptional regulators that promote PB differentiation and function (*IRF4*, *ELL2*, and *XBP1*) as well as chemokine receptors that are normally down-regulated for proper positioning and clustering of GCBCs (*GPR183* (EBI2) and *CCR7*)^26^ (**Supplementary Table 2**). This BCL6- dependency of the preGC transcriptional program, analyzed at single-cell resolution, provides important supporting evidence that the *in vitro* preGC trajectory recapitulates a key aspect of the transcriptional requirement for BCL6 that underlies GCBC initiation *in vivo*.

### Temporal profiling of chromatin and gene expression dynamics in bifurcating B cell trajectories

To assemble high-resolution GRNs underlying bifurcating B cell fate trajectories, we performed temporal single-cell multiomics, jointly profiling gene expression and chromatin accessibility at 24h intervals. For these analyses, we used two different donors (male and female). To resolve discrete B cell states, we performed MIRA-based joint topic modeling, representing each cell as a weighted mixture of gene expression and chromatin accessibility topics (i.e., groups of genes or peaks that co-vary across cells)^12^ (**Fig. 2a** and **Extended Data Fig. 2a**). The analysis revealed 27 expression topics and 13 accessibility topics that were specifically associated with at least one B cell state (**Extended Data Fig. 2b**). Most of the expression topic gene sets were significantly overlapping between the two different donors (**Extended Data Fig. 2c**). Furthermore, the majority of accessibility topics contained ATAC-seq peaks that were linked to sets of genes that significantly overlapped those of corresponding gene topics, consistent with the expectation that the accessibility topics contain regulatory elements that control transcription of genes in expression topics in a cell state-specific manner (**Extended Data Fig. 2d**). As with the scRNA-seq dataset (**Fig. 1c**), SVM training on *in vivo* B cell states derived from the Human Tonsil Atlas^14^ was used to classify each cell in our multiome datasets (**Fig. 2a**). Cells from the D0-D2 timepoints predominantly had high probabilities of naïve or ActB states. The probability of GCBC and PB classification increased at D3-4 timepoints as the ActB probability decreased. Finally, in agreement with the scRNA-seq analysis (**Fig. 1d**), D5-D6 cells were predominantly classified as GCBC or PB. Leiden clustering identified eight B cell states annotated as naïve, ActB-1 to ActB-4 (corresponding to D1 to D4), preGC-1, preGC-2, PB-1 and PB-2 (**Fig. 2a** and **Extended Data Fig. 2a**).

**Fig. 2.**
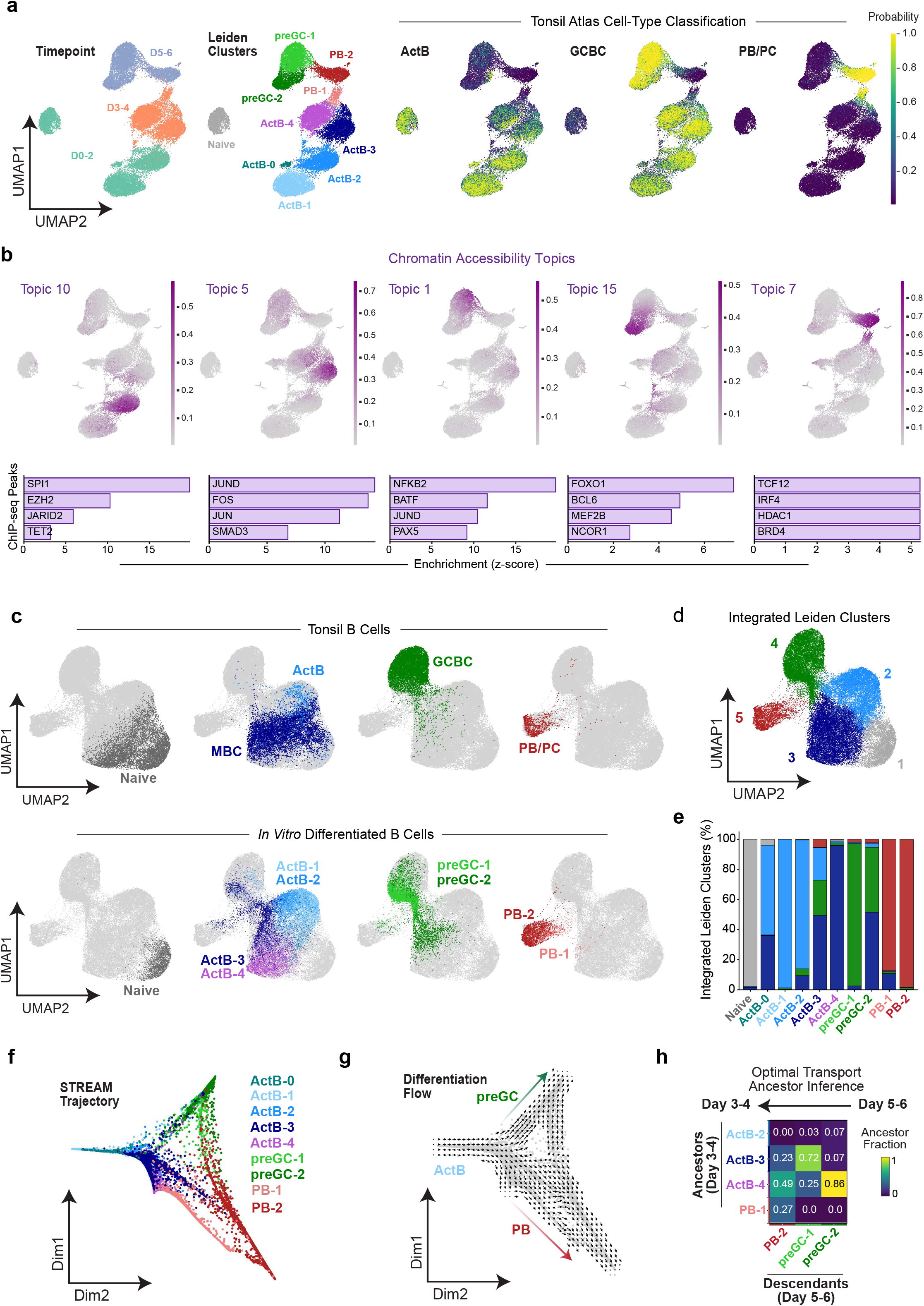
Temporal profiling of chromatin and gene expression dynamics in bifurcating B cell trajectories. **(a)** Naïve human B cells from Donor 1 (male) were activated and differentiated as in Fig. 1a and profiled every 24 hours by paired single-cell RNA/ATAC-seq. MIRA joint topic modeling of transcripts and chromatin accessibility features was used to jointly project cells in low-dimensional UMAP space and colored based on timepoint (*left*), Leiden clusters with their cell-type annotations (*middle*), and support vector machine classification probabilities based on Human Tonsil Atlas cell-types (*right*). **(b)** UMAP projections of chromatin accessibility topic composition (purple) are displayed for representative ActB-associated (topic 10 and topic 5), preGC-associated (topic 1 and topic 15), and PB-associated (topic 7) topics. ChIP-seq peak enrichment for the top 10,000 peaks is displayed beneath each indicated accessibility topic. **(c)** Harmony-integrated UMAP of tonsil and *in vitro* differentiated B cells, colored by annotations from Human Tonsil Atlas (*top*) or Fig. 2a (*bottom*). **(d)** UMAP displaying cells colored according to Leiden clustering of the integrated Harmony object. **(e)** Stacked bar plot displaying distributions of *in vitro* differentiated B cells in Leiden clusters from Fig. 2d. **(f)** STREAM trajectory analysis of *in vitro* differentiated B cells, colored by their annotations from Fig. 2a. **(g)** STREAM pseudotime values were used with the CellOracle gradient module to generate differentiation flow vector fields (see Methods). **(h)** Moscot optimal transport analysis of ancestral relationships. Heatmap showing backward transition probabilities from Day 5–6 descendant states (columns) to Day 3–4 ancestor states (rows).

Gene ontology analysis revealed that the expression topic gene sets were enriched for a wide range of important immune-cell pathways associated with GC biology or extrafollicular PB responses (**Supplementary Table 3**). ChIP-seq peak enrichment analysis using over six thousand uniformly processed datasets from the Cistrome Data Browser (CistromeDB)^27^ revealed that the accessibility topic peak sets were enriched for regions bound by known regulators of activated B cell, GCBC, or PB transcriptional programs (**Fig. 2b** and **Supplementary Table 4**). For example, ActB-associated topic 10 was enriched for regions bound by SPI1 (PU.1) and repressive epigenetic modifiers (EZH2, JARID2, TET), suggesting these regions may become de-repressed following activation. In contrast, ActB-associated topic 1 was enriched for regions bound by AP-1 family members. AP-1 binding regions were also enriched in preGC-associated topic 5 along with those for NFKB1. As expected, one of the preGC-associated topics (#15) was enriched for regions bound by FOXO1 and BCL6 and its co-regulators MEF2B and NCOR1. In contrast, the PB-associated topic was dominated by enrichment for IRF4, TCF12 and chromatin remodelers (HDAC1 and BRD4).

To further analyze the correspondence of *in vitro* B cell states to the continuum of *in vivo* B cell states, we integrated the gene expression profiles of our *in vitro* B cells with those in the Human Tonsil Atlas using Harmony^28^. Following dimensionality reduction, the Harmony-integrated data set revealed five major clusters corresponding to naïve, ActB, PB and GCBC states with contributions from the Tonsil Atlas B cell dataset (*top*) as well as the B cell states generated by our *in vitro* system (*bottom*) (**Fig. 2c,d**). Notably, the *in vitro* generated preGC cells appeared to bridge the tonsil ActB and GCBC states, consistent with their identity as a precursor state that is poised for GCBC differentiation. Furthermore, in contrast to the *in vitro* ActB clusters, cells in the preGC clusters were predominantly part of the integrated GCBC cluster where tonsil GCBCs were largely positioned (integrated Leiden cluster 4) (**Fig. 2d,e**).

To rigorously infer trajectory bifurcation, we utilized the STREAM pipeline, which infers pseudotime values and differentiation branch points based on variable gene expression patterns^29^ (**Fig. 2f** and **Extended Data Fig. 2e**). These pseudotime values were then used in the CellOracle modeling framework to generate a differentiation gradient which estimates the flows of differentiation that can be represented as a vector field graph^30^ (**Fig. 2g** and **Extended Data Fig. 2f**). The analysis revealed that the majority of annotated PB and preGC cells for each donor dataset resided in divergent trajectories emanating from a common branchpoint comprising ActB cells. We additionally applied MultiVI dimensionality reduction and multi-modal optimal transport (OT) analysis framework, Moscot^31^, that jointly leverages chromatin accessibility and transcriptome to estimate likely transitions between cell states. Importantly, the MultiVI train-validation reconstruction loss gap remained stable (5.6% of validation loss at epoch 10 versus 6.1% at epoch 200), indicating that the latent space did not become fit to noise in the training cells (**Extended Data Fig. 2g**). We computed the backward transition (pull-back distribution) probabilities from the Day 5-6 cell states (descendants) to Day 3-4 cell states (ancestors) (**Fig. 2h**). These results suggested that while the final PB state (PB-2) descends from progenitors emanating from the ActB-3, ActB-4 or PB-1 cell states, preGC-1 and -2 descend exclusively from the ActB-3 and -4 cell states.

Collectively, the temporally resolved multiomics profiling demonstrates that *in vitro* activated human B cells bifurcate into distinct trajectories reflecting key transcriptional and chromatin features of *in vivo* PB and GCBC states. Importantly, the *in vitro* generated preGC cells occupy an intermediate state between activated B cells and tonsil GCBCs, in keeping with the evidence in Fig. 1 that they represent a poised GC precursor state. Therefore, the *in vitro* system and the temporally resolved multiomic profiling provide a tractable system to comprehensively investigate the GRNs that govern PB versus preGC fate choice in activated human B cells.

### Assembly of GRNs underlying bifurcation of human B cells into PB and preGC fates

We sought to comprehensively assemble B cell state-specific GRN models spanning multiple scales linking TFs, candidate cis-regulatory elements (cCREs), target genes and fate choice. To do so, we integrated two complementary modeling frameworks previously used independently: MIRA regulatory potential modeling^12^ and CellOracle GRN inference^30^. The following considerations motivated this design. Traditional GRN models infer TF-gene linkages by assessing TF motif enrichment at candidate cis-regulatory elements (cCREs) that are co-accessible with gene promoters^30,32^. This approach has two limitations: (1) gene expression can be dynamically regulated independently of alterations in cCREs and promoter accessibility and (2) a given TF motif can be bound by structurally related or even unrelated TFs. To overcome these two limitations, the first phase of our GRN assembly utilized MIRA regulatory potential (RP) modeling^12^ to establish binary TF-gene linkages (**Fig. 3a**). Instead of relying on promoter-and- cCRE accessibility correlations, MIRA takes advantage of single-cell multiome data to construct RP models for each dynamically expressed gene based on the correlation between accessibility of associated cCREs and gene transcripts across cells. In RP modeling, the influence of a cCRE on gene expression is additive and decays exponentially with genomic distance from the transcription-start site (TSS), an assumption supported by chromatin-conformation data such as Hi-C^33^. Importantly, MIRA leverages over ten thousand uniformly processed ChIP-seq datasets in CistromeDB rather than TF motif enrichment to infer binding of annotated TFs to cCREs. Probabilistic in silico deletion (ISD) of TF binding regions then generates gene-specific models predicting TF-gene regulatory relationships based on how removal of TF binding peaks affects the model’s ability to predict transcript levels from accessibility features. This approach has successfully been applied to robustly predict TF-gene linkages^34,35^. Thus, in Phase I, the MIRA assembled GRN is based on dynamic chromatin and gene expression profiles of human B cells and their integration with evidence of physical TF-cCRE interactions based on CistromeDB ChIP-seq datasets, providing a robust set of TF-gene linkages for downstream network refinement.

**Fig. 3.**
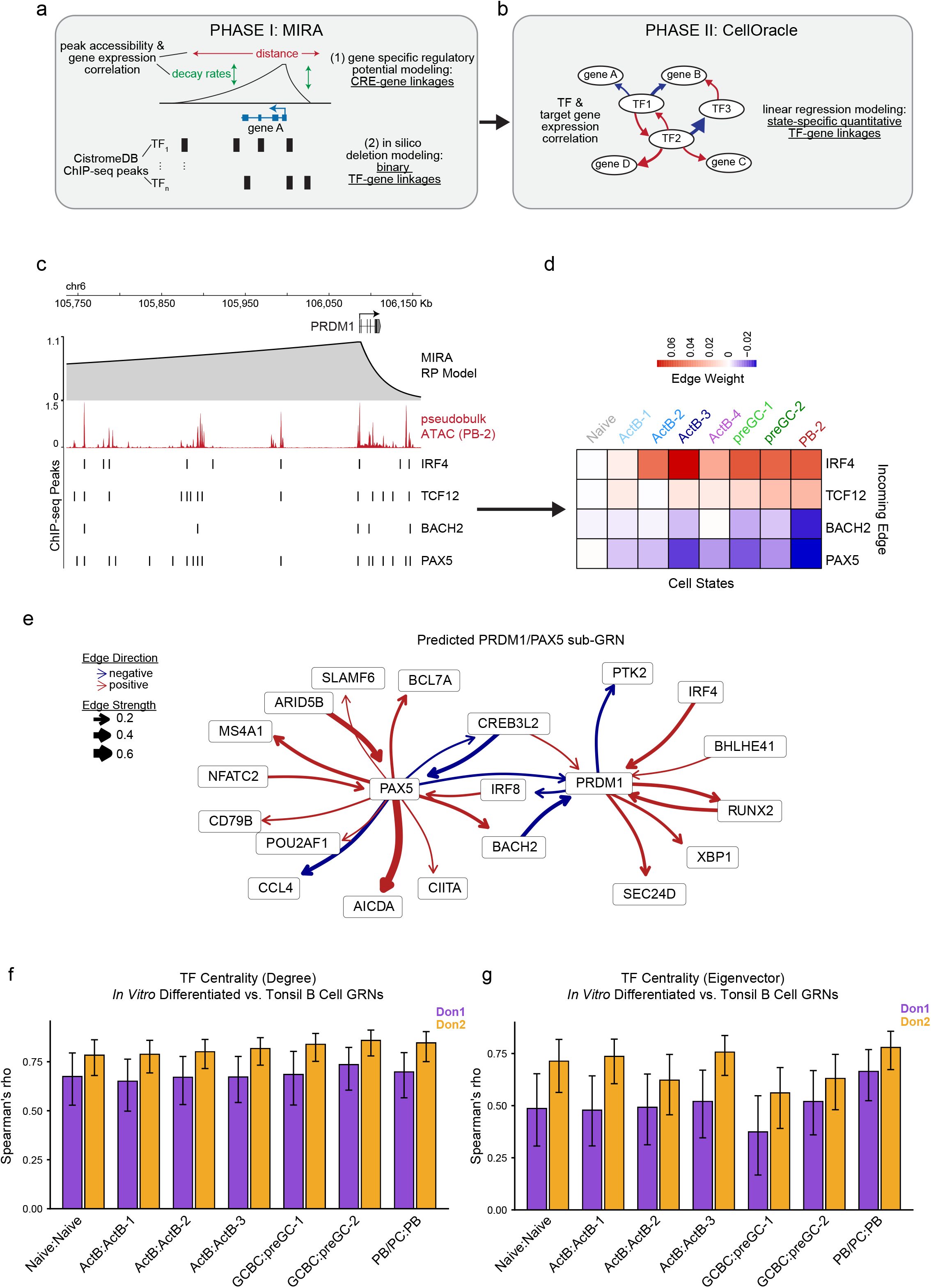
Assembly of GRNs underlying bifurcation of human B cells into PB and preGC fates. **(a)** Phase I schematic: assembly of qualitative TF-gene linkages using MIRA regulatory potential (RP) and in silico deletion (ISD) modeling. Gene-specific RP models incorporate learned decay rates and distances based on correlations between peak accessibility and gene expression from multiome data. TF regulatory potential was inferred by measuring RP model changes following probabilistic ISD of TF binding regions (CistromeDB ChIP-seq peaks). **(b)** Phase II schematic: CellOracle transforms binary MIRA-derived TF-gene linkages into quantitative, directional, state- specific linkages using Bayesian ridge regression. **(c)** The RP model for *PRDM1* is shown, with the regulatory potential (gray shading) assigned to flanking ATAC-seq peaks as a function of distance and learned decay rates from the TSS. ATAC-seq signals derived from the PB state pseudobulk are shown below the RP model. CistromeDB ChIP-seq peaks that overlap with ATAC- seq peaks for representative TF regulators predicted by ISD modeling are shown below. **(d)** Representative CellOracle state-specific TF-gene edge weights for MIRA-predicted regulators of *PRDM1*. **(e)** Representative predicted sub-GRN for PAX5 and PRDM1 derived from CellOracle GRN modeling. Select incoming and outgoing edges for the PAX5/PRDM1 nodes in the PB state are displayed with thickness of arrows corresponding to state-specific edge weight values. Red and blue edges indicate positive and negative regulation, respectively. (**f**, **g**) Comparison of *in vitro*-derived and tonsil B cell GRN TF centrality. Bar charts display the Spearman’s rank correlation between degree (**f**) and eigenvector (**g**) centrality scores for TFs in the indicated tonsil- *in vitro* B cell state pairs. Comparison was restricted to TFs that were present in both GRNs for each comparison. In vitro-derived GRNs from Donor 1 (male) and Donor 2 (female) are indicated by purple and orange bars, respectively.

In the second phase of GRN assembly (**Fig. 3b**), we used the binary TF-gene linkages derived from MIRA as a base-GRN that serves as input for inference of context-specific GRNs using CellOracle^30^. As we experimentally test and demonstrate later, this base-GRN is of higher quality than the default base-GRN of CellOracle which is based on promoter-peak co-accessibility and TF motif enrichment. CellOracle infers cell state-specific GRNs using Bayesian ridge regression to fit TF and target gene transcript levels, estimating TF contributions to target gene activity within each state. This enables TF-gene linkages to be assigned quantitative edge weights (strengths) as well as signs (activation or repression). These transformed TF-gene linkages can then be used to infer the centrality of each TF to a given cell state and predict the change in cell states using *in silico* TF perturbation simulations. A methodological advance is the coupling of MIRA and CellOracle that significantly enhances the quality of the base-GRN and makes the false discovery rate (FDR) control of inferred regulatory relationships more stringent (see testing in **Fig. 4e** and **Extended Data Fig. 6d**).

**Fig. 4.**
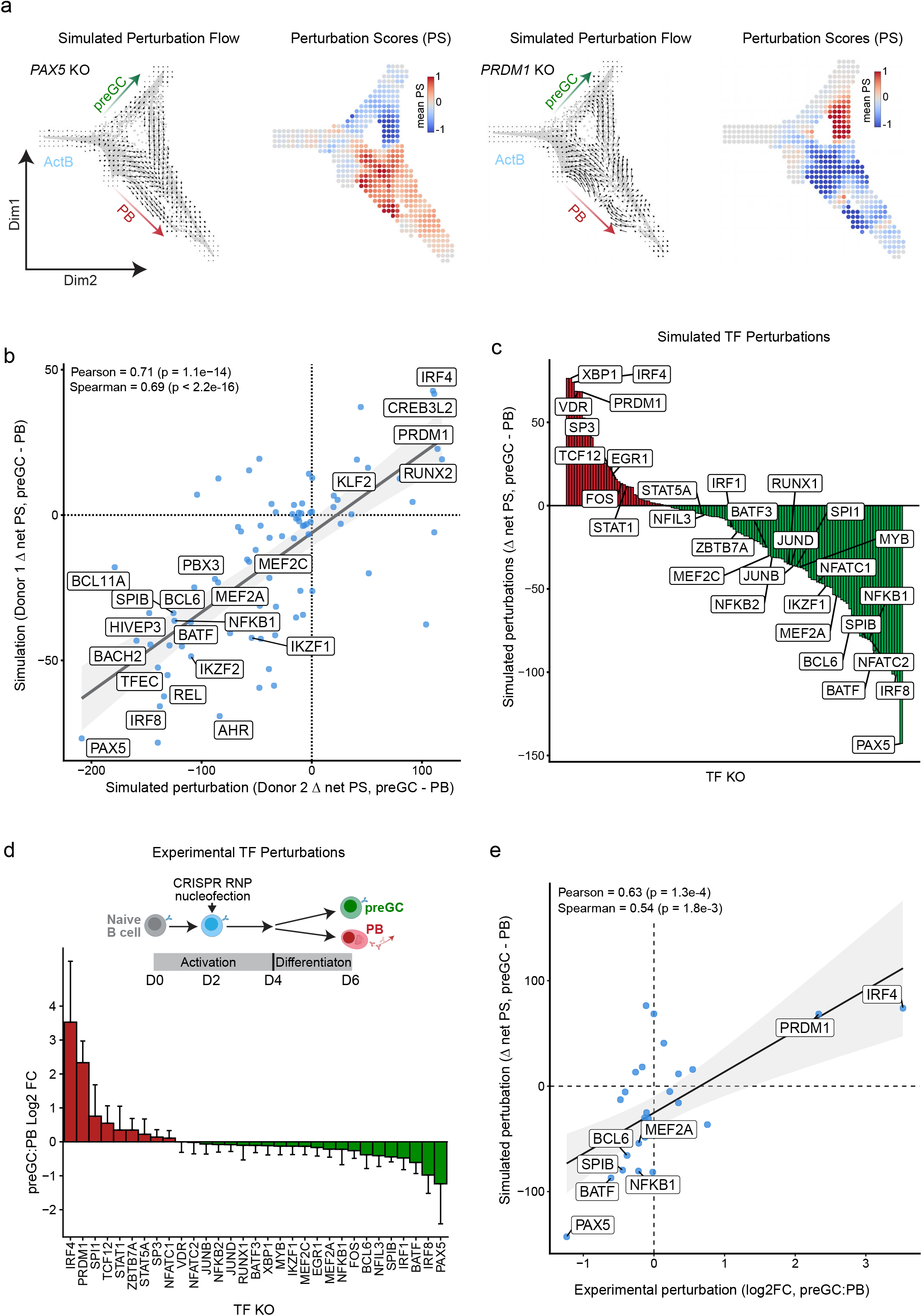
Predicting and testing TF perturbation effects on preGC versus PB trajectories. (**a**) Representative *in silico* TF KO simulations visualized with perturbation flow vector fields or perturbation score grids for PAX5 (*left*) or PRDM1 (*right*) (Donor 1). Perturbation scores (PS) are the dot product of differentiation flow vectors and perturbation simulation vectors. Net preGC and net PB perturbation scores (PS) are derived from summing the PS values for cells in preGC or PB states, respectively. **(b, c)** Delta net perturbation scores (preGC PS minus PB PS, KO vs. control) for 124 TF KO simulations. (**b**) Correlation between Donor 1 and Donor 2 TF KO simulation results. Scatter plot displays the delta net PS for each TF for each donor, with linear regression fit (gray line) and 95% confidence interval (shaded band). (**c**) Bar plot displays mean delta net PS for each TF, ranked in descending order. Positive values indicate pro-PB TFs; negative values indicate pro-preGC TFs. **(d)** Experimental results of arrayed CRISPR TF KO screen. Naïve B cells were activated for 2 days before nucleofection with CRISPR RNP complexes and re-culture as in Fig. 1a. Cells were harvested at D6 for preGC (IRF8^hi^IRF4^lo^) and PB (IRF8^lo^IRF4^hi^) quantification by flow cytometry (n = 4 experimental replicates). Bar plot displays log2 fold change in the preGC:PB ratio (D6) of KO vs. NTC (non-targeting control) for indicated TFs (n = 4-7, 2-4 experimental replicates each for Donor 1 and Donor 2). (**e**) Comparison of simulated and experimental TF KO effects on B cell fates. Scatter plot displays mean simulated perturbation scores (net preGC PS - net PB PS) and the experimentally determined mean log2 fold change in preGC:PB ratios (KO vs. control) for the 31 TFs selected for experimental testing (see Fig. 4c). Gray line and shaded band indicate linear regression fit and 95% confidence interval, respectively.

To illustrate Phase I, we show the MIRA RP model for the PB-associated gene *PRDM1* (**Fig. 3c**) as an example. The relative importance of each cCRE is weighted as a function of bi- directional learned decay rates and distances from the gene’s TSS. Notably, the gene-specific RP models generally captured the TSS-cCRE interactions that were detected by chromatin conformation capture assays of accessible regions (HiCAR) in the GM12878 human B cell line^36^ (see UCSC Genome Browser session: https://tinyurl.com/BcellMultiomeTracks). The cCREs in these gene-specific RP models were annotated with CistromeDB ChIP-seq TF binding datasets (selected TF ChIP-seq peaks are displayed below ATAC-seq peaks in **Fig. 3c**). This in turn enabled prediction of the regulatory impact of a given TF on its target gene by probabilistic *in silico* deletion (ISD) modeling (see Methods). The top 5% of TF-gene ISD scores were then used to generate binary TF-gene linkages that served as the base-GRN for inference of context-specific GRNs using CellOracle.

In Phase II, CellOracle transformed the binary TF-gene linkage predictions of MIRA into quantitative TF-gene edge strengths for each B cell state (**Supplementary Table 5**). For example, CellOracle predicted IRF4 and TCF12 as positive regulators of the *PRDM1* gene and PAX5 and BACH2 as negative regulators, each with dynamic edge strengths across the various B cell states (**Fig. 3d**). To generate the highest-confidence, state-specific GRNs, we refined the initial GRNs by re-fitting each model with the top 10,000 statistically significant TF-gene edges (p < 0.001). This integrated analysis provides a catalogue of predicted human B cell state-specific TF-gene linkages, which are highly dynamic and undergo pronounced reconfigurations during B cell activation and divergence into PB and preGC trajectories (**Supplementary Table 5**). For example, we show the predicted PRDM1/PAX5 sub-GRNs that recapitulate several cross-regulatory relationships well established in the murine system (**Fig. 3e**). To facilitate the accessibility and use of these GRN predictions, we have generated a programming-free, interactive web application (https://pitt-csi.shinyapps.io/humanBcellGRN/).

We next sought to determine if applying the same MIRA + CellOracle framework to primary human B cells would generate similar state-specific GRN models. Thus, we used single-cell multiomics data from the Human Tonsil Atlas to generate GRNs for the five major B cell states (Naïve, ActB, MBC, GCBC, and PB/PC; see Fig. 2c). We first compared the chromatin accessibility profiles of the corresponding differentiated cell states (*in vitro* preGC vs. tonsil GCBC and *in vitro* PB vs. tonsil PB/PC) by intersecting ATAC-seq peaks called from the respective pseudobulk clusters. We found considerable overlap, with Jaccard indices reaching 0.40 (*in vitro* preGC vs. tonsil GCBC) and 0.45 (*in vitro* PB vs. tonsil PB/PC) at the most stringent peak threshold (q < 1e-5) (**Extended Data Fig. 3a,b**). At a more permissive threshold (q < 1e-2), the number of overlapping peaks increased to ∼100,000 (preGC vs. GCBC) and ∼115,000 (PB vs. PB/PC). Notably, a subset of peaks at the PRDM1 locus was detected only in the *in vitro* PB and tonsil PB/PC states and not in *in vitro* preGC or tonsil GCBC states, demonstrating correspondence of state-specific accessibility features between *in vitro* and *in vivo* generated B cell states for a key regulator (**Extended Data Fig. 3c,** see https://tinyurl.com/BcellMultiomeTracks for genome-wide views of *in vitro* and tonsil B cell ATAC-seq tracks).

After applying the MIRA + CellOracle framework to the tonsil B cell multiome dataset, we compared the resulting GRN architectures to those of the *in vitro* differentiated B cells. We found that the centrality of shared TFs was significantly correlated between corresponding *in vitro* vs. tonsil B cell state pairs, with Spearman’s correlation coefficients reaching 0.78 for degree centrality and 0.75 for eigenvector centrality scores (**Figs. 3f,g** and **Supplementary Table 6**). The degree centrality tended to be more strongly correlated than eigenvector centrality, suggesting the coarse-grained network topologies are well conserved while finer influence- weighting is more variable between the *in vitro* vs. tonsil B cells. At the level of individual TF regulons (i.e., target genes), TF-gene edge-weight rank correlations were significant across all state pairs, with median Spearman’s correlation coefficients of 0.25–0.50 (**Extended Data Fig. 3d** and **Supplementary Table 5**).

To evaluate the quality of individual TF regulon predictions, we compared BCL6 target gene predictions with the experimentally validated target genes from the BCL6-KO scRNA-seq experiment (**Fig. 1h**). As expected, given that the experimental data were derived from *in vitro* differentiated preGCs, the *in vitro* preGC GRN models showed the greatest concordance between observed and predicted target genes, with odds ratios (OR) of ∼5 to 6 (**Extended Data Fig. 3e**). The overlapping observed and predicted target genes included known BCL6 target genes (e.g. *BCL2*, *CCND2*, *MIR155HG*) as well as previously unreported target genes that are associated with immunoglobulin production (*ELL2*) and the endoplasmic reticulum/unfolded protein response (*SSR4*, *PPIB*, *HSP90B1* (GRP94)). Nonetheless, the predicted BCL6 repressed target genes in the tonsil GCBC GRN also showed significant enrichment for BCL6-KO up-DEGs (i.e., experimentally repressed genes) (OR = ∼2; p = 0.04). Together with the substantial overlap between *in vitro* and tonsil B cell GRNs, these results indicate that the GRN modeling framework generates TF-gene predictions that are (1) conserved between *in vitro* and tonsil B cells and (2) enriched for experimentally validated BCL6 target genes.

### Predicting and testing TF perturbation effects on preGC versus PB trajectories

The assembly of human B cell state-specific GRN models enabled comprehensive predictions of the impact of individual TFs on preGC/GCBC versus PB/PC trajectories by simulating their perturbations *in silico*. CellOracle assessed such perturbations by propagating the loss of expression of a given TF across the inferred GRN via its primary, secondary and tertiary target genes. The resulting changes in gene expression values were used to estimate shifts in cell states which can be used to calculate and visualize perturbation flow vector graph fields projected onto the STREAM trajectories. For example, simulation of the *PRDM1* KO results in a change in differentiation flow away from the PB trajectory and towards the preGC trajectory while simulation of *PAX5* KO produces the opposite pattern (**Fig. 4a**), consistent with their well-established roles in promoting or antagonizing PB differentiation, respectively. These simulated perturbation flows were well conserved both for the *in vitro* B cell Donor 2 GRN (**Extended Data Fig. 4a**) and for the tonsil B cell GRN (**Extended Data Fig. 4b-d**). For each TF KO simulation, a perturbation score was generated for the GC-associated trajectory (*in vitro* preGC, tonsil GCBC) and the PB-associated trajectory (in vitro PB, tonsil PB/PC) by calculating the dot product of differentiation flow vectors and perturbation flow vectors. To systematically predict the effect of central TFs and nominate new regulators of the bifurcating trajectories, we filtered on genes that were classified as TFs^37^ with an eigenvector centrality greater than 0.005 (**Supplementary Table 6**). Of these, 124 TFs were represented as source and target nodes within the final GRN models and thus could be simulated with *in silico* knockout modeling. The resulting simulations using the *in vitro* Donor 1 GRN produced a wide range of predicted effects on the preGC versus PB differentiation that were strongly correlated with those observed using the *in vitro* Donor 2 GRN, suggesting the predictions are robust across experiments, donors and sex (**Fig. 4b, Supplementary Table 7**). In addition to correctly predicting the effects of well-established pro- PB regulators (e.g. PRDM1, IRF4) and pro-GC regulators (e.g. PAX5, BACH2, BCL6, IRF8, BATF, NFKB1, MEF2C), these models also nominated less-appreciated regulators of the PB (RUNX2, CREB3L2, KLF2) and preGC (BCL11A, MEF2A, TFEC, IKZF1/2, PBX3, HIVEP3) trajectories. We next evaluated how well these predictions correlated with those generated using the tonsil B cell GRN model. This revealed a significant correlation between the *in vitro* and tonsil B cell GRN predicted effects on each pair of bifurcated trajectories (Pearson’s correlation, r = 0.30, p = 6.5e-3; Spearman’s correlation, rho = 0.35, p = 1.9e-3) (Extended Data Fig. 4e). While the magnitude of this global correlation was modest, the concordance was driven by directionally consistent recovery of the same key regulators in both settings. Importantly, many of the novel candidate TF regulators were preserved (PB/PC: KLF2, CREB3L2, RUNX2; GCBC: MEF2A, TFEC, IKZF1/2, PBX3, HIVEP3), suggesting these previously unappreciated TFs may play important roles in B cell fate dynamics *in vivo*.

To comprehensively assess the quality of the GRN models and the resulting *in silico* perturbation predictions, we used an arrayed CRISPR screen to knockout the corresponding TF genes and assay their impact on the preGC versus PB trajectories using the *in vitro* system. We selected 31 TFs that (1) had documented roles in regulating murine B cell differentiation or function and (2) showed a wide range of predicted effects on the bifurcating trajectories in our simulations (**Fig. 4c** and **Supplementary Table 8**). For experimental testing of these predictions, naïve B cells for each donor were separately activated for two days before nucleofection with CRISPR RNP complexes containing gRNAs targeting a selected TF gene (**Fig. 4d**). The cells were then cultured for four additional days as detailed in Fig. 1. PreGC (IRF8^hi^IRF4^lo^) and PB (IRF8^lo^IRF4^hi^) cell numbers at Day 6 were quantitated by flow cytometry (see Fig. 1a for example gating strategy). CRISPR editing efficiency was confirmed by extracting DNA from cells on Day 4 for PCR genotyping (**Extended Data Fig. 4f**). The results of the experimental screen, averaged across 4-7 replicates with the male and female donors, revealed 7 TF KOs that decreased the mean preGC:PB ratio by >20% (PAX5, IRF8, BATF, IRF1, SPIB, NFIL3, BCL6) and 6 TF KOs that increased the mean preGC:PB ratio by >20% (IRF4, PRDM1, SPI1, TCF12, STAT1, ZBTB7A) (**Fig. 4d** and **Supplementary Table 8**). Of these, only PRDM1, IRF4, IRF8, BATF and SPIB had q-values < 0.1, consistent with these being central regulators of the preGC and PB trajectories as previously reported in murine experiments. Importantly, the simulated and experimental preGC:PB ratios resulting from the TF perturbations were highly correlated (Pearson’s correlation, r = 0.63, p = 1e-4; Spearman’s correlation, rho = 0.54, p = 2e-3) (**Fig. 4e**). This correlation was stronger than one generated by comparing the experimental results with simulations performed using the default Cicero base-GRN input into CellOracle (Pearson’s correlation, r = 0.51, p = 1e-3; Spearman’s correlation, rho = 0.31, p = 9e-2; **Supplementary Table 8**). The improved performance achieved by coupling MIRA and CellOracle demonstrates the value of integrating these complementary frameworks for GRN assembly. Collectively, the results represent an important test of the assembled human B cell GRN models.

### ChromBPNet uncovers dominant and reciprocal TF action at IRF motifs during B cell fate specification

While MIRA-CellOracle integration enabled assembly of B cell state-specific GRNs with quantitative and directional TF-gene linkages, these models did not include individual state-specific TF-CRE linkages that represent state-specific TF action at binding sites within CREs. To address this gap, we applied ChromBPNet to infer B cell state-specific TF activity across all ATAC-seq peaks at base-pair resolution. ChromBPNet is a convolutional neural network (CNN) that learns the contribution of individual base-pairs to the accessibility of ATAC-seq peaks within which they are embedded, after regressing out Tn5 enzyme cutting bias^38^. We applied this CNN framework to our multiome dataset by training state-specific ChromBPNet models on pseudobulk ATAC-seq reads from each cell cluster. To ensure that each model was not being overfitted to the data, testing was performed using held-out chromosome peaks (see **Supplementary Table 9** for ChromBPNet bias model evaluation metrics). Each of these state-specific models was then used to generate corresponding base-pair resolution contribution score (CS) predictions across the union set of ATAC-seq peaks (**Fig. 5a**). These state-specific contribution score tracks revealed short DNA sequences, referred to as “seqlets”, that were predicted to be important in determining the accessibility profiles of their corresponding ATAC-seq peaks. Strikingly, many of the ChromBPNet seqlets corresponded to known TF motifs, thereby implicating action of cognate TFs in binding to such sequences and controlling chromatin accessibility. Examples of seqlets contributing to the accessibility of cCREs in the *AICDA* and *PRDM1* loci in the preGC-2 and PB cell states, respectively, are shown in **Fig. 5b**. The former represents the motif for the AP-1 family of TFs, whereas the latter is bound by the POU domain family members Oct-1,2.

**Fig. 5.**
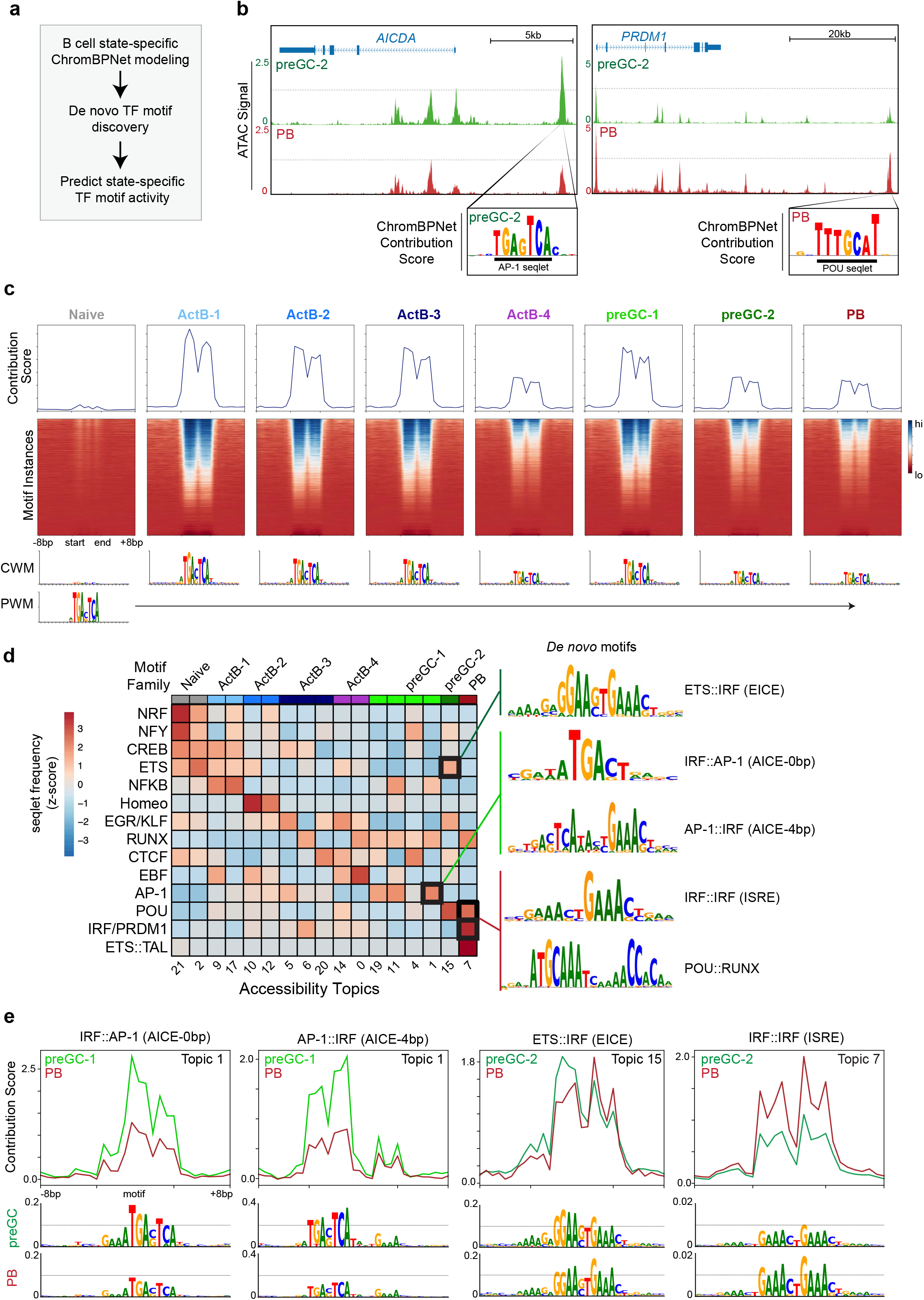
ChromBPNet uncovers dominant and reciprocal TF action at IRF motifs during B cell fate specification. **(a)** Workflow for B cell state-specific ChromBPNet modeling: training state-specific models, de novo TF motif discovery, and prediction of state-specific TF activity. **(b)** UCSC genome browser views of AICDA (*left*) and PRDM1 (*right*) loci showing ATAC-seq tracks in preGC-2 and PB states with ChromBPNet contribution scores. Seqlets corresponding to AP-1 (AICDA) and POU (PRDM1) motifs are highlighted. **(c)** Aggregated ChromBPNet contribution scores at all AP-1 motifs in union set of ATAC-seq peaks across 8 B cell states (*top*). DNA sequences aligned on AP-1 motif center (7 bp) ± 8 bp flanking. Base-pair resolution contribution score heatmaps (*middle*) with motif instances (rows) ranked by CWM-PWM dot product. State-specific CWM and reference PWM logos shown below. **(d)** TFModisco-derived seqlet frequencies (z-scaled) for TF motif families across accessibility topics (*left*). Analysis performed on top 10,000 peaks per topic using state-specific ChromBPNet models; seqlets annotated with JASPAR 2024 motif clusters. Only families with ≥5% frequency in at least one topic are shown. CWMs for dominant IRF-containing composite elements in preGC (EICE, AICE) versus PB (ISRE) states are shown (right). **(e)** Aggregated contribution score signals (*top*) and CWM logos (bottom) for preGC versus PB states at topic peaks containing IRF-related composite elements (AICE, EICE) or ISRE motifs.

Conventional TF motif analysis tools (e.g., HOMER) are inherently restricted to reporting nucleotide enrichment relative to a background set, with the resulting position weight matrix (PWM) being a fixed summary of sequence composition at a set of aligned instances that cannot vary with cellular context. In contrast, sequence-to-function models like ChromBPNet that are trained on cell state-specific genomic features can report how much each nucleotide contributes to that feature, with the resulting contribution score weight matrix (CWM) composed of base-pair weights that vary with cellular context. Therefore, we used the outputs of the eight B cell state-specific ChromBPNet models to analyze, in a directed as well as unbiased manner, the dynamical actions of TF families on all cCREs containing their cognate TF motifs. To illustrate the directed analysis, we displayed aggregated base-pair contribution score (CS) profiles at all instances of the AP-1 motif across the union set of ATAC-seq peaks encompassing the distinct B cell states (**Fig. 5c**, *top*). The aggregated CS profiles of the AP-1 motif-containing cCREs were highly dynamic across the various B cell states, with pronounced biphasic activity that peaked in the ActB-1 and preGC-1 states as indicated by changes in contribution weight matrix (CWM) logo heights (**Fig. 5c**, *bottom*). The former is consistent with the rapid activation of AP-1 activity downstream of BCR and CD40 signaling^39,40^ and the latter is consistent with activity in response to restimulation with CD40 signaling that promotes the preGC trajectory. To analyze AP-1 activity at a given DNA region containing the motif, we used dot product calculations to quantify the similarity between the CWM of each motif instance and a reference position weight matrix (PWM) from Jaspar (MA1634.2) (see Methods). We then hierarchically arrayed the AP-1 motif instances according to their CS dot products in a reference B cell state (ActB-1) enabling visualization of the base-pair contribution scores for each motif instance across all states (**Fig. 5c**, *middle*). This revealed a wide range of predicted contributions of AP-1 motifs to chromatin accessibility within each state. Thus, these B cell state-specific ChromBPNet models establish a compendium of B cell state-specific putative TF-cCRE regulatory linkages that represent a critical layer of the multi-scale GRN assembly.

To evaluate dominant TFs regulating the state-specific chromatin accessibility programs in an unbiased manner, we used TFModisco to analyze the ChromBPNet contribution scores within the top 10,000 ATAC-seq peaks of each state-specific accessibility topic delineated by MIRA (**Extended Data Fig. 2b**). TFModisco is a motif discovery algorithm that first identifies seqlets and performs iterative meta-clustering to aggregate seqlets with similar contribution score weight matrices to generate *de novo* motifs^41^. We note that *de novo* motif discovery can also be performed using conventional, frequency-based tools such as HOMER; however, we sought to investigate predictions made from the B cell state-specific ChromBPNet models given the added benefit of inferring state-specific base-pair contributions to accessibility rather than base-pair frequency. In the final step, the best match for each *de novo* motif pattern is determined by comparing it to known Jaspar TF motifs. We additionally merged related motif hits in each motif family (see **Supplementary Table 10**) and then quantified the motif family seqlet frequencies across the various chromatin accessibility topics (see representative TF family *de novo* motifs in **Extended Data Fig. 5a**). This enabled unbiased and comprehensive delineation of TF families acting on cCREs that were reflective of specific accessibility topics and B cell states (**Fig. 5d** and **Extended Data Fig. 5b**). Since many different TFs within a family can recognize the same motif, we also analyzed the transcript levels of all cognate TFs within each motif family (**Extended Data Fig. 5c** and **Supplementary Table 11**). The combined analysis provided fine-resolution predictions for which TFs are most likely responsible for driving the dynamic chromatin programs during human B cell activation and differentiation. Importantly, these orthogonally derived state-specific TF activity predictions closely aligned with corresponding TF GRN centrality scores (**Supplementary Table 7**).

Strikingly, the TFModisco analysis, based solely on state-specific contribution score tracks, identified de novo motifs that matched IRF4/IRF8 composite elements (CEs) within the preGC- and PB-associated topics (**Fig. 5d**). Specifically, the preGC-1 topics contained AP-1::IRF (AICE) CWM motifs which are cooperatively bound by BATF and IRF4/IRF8, and the preGC-2 topics contained ETS::IRF (EICE) CWM motifs which are cooperatively bound by PU.1/SPIB and IRF4/IRF8. In contrast, the PB topic was enriched for IRF::IRF (ISRE) CWM motifs which are bound by IRF4/IRF8 homo- or heterodimers as well as PRDM1^5,42^. Importantly, in addition to recovering known IRF-related composite elements, this analysis also uncovered a novel putative composite element composed of POU and RUNX motifs. While RUNX2 and Oct-1 (*POU2F1*) have been reported to form complexes on juxtaposed motifs at the β-casein promoter^43^, the POU::RUNX CE configuration identified here has not been previously described. Interestingly, the POU::RUNX CE seqlets were enriched in the PB topic (#7), suggesting they may play a critical role in establishing or maintaining PB chromatin states.

We next used a composite element motif scanning tool developed by us, CEseek^44^, to identify all instances of each of the aforementioned CEs within their respective accessibility topics and compared the contribution score signals at these regions between cell states. As expected, the AICE-containing preGC-1 topic (#1) peaks were more highly accessible in the preGC-1 state compared to the PB state and EICE-containing preGC-2 topic (#15) peaks were more accessible in the preGC-2 state compared to the PB state (**Extended Data Fig. 5d**). Conversely, the ISRE-containing PB topic (#7) peaks were more accessible in the PB state compared to the preGC states. The aggregated contribution scores revealed substantial increases in the predicted contribution at AICE motifs in the preGC-1 state and ISRE motifs in the PB state (**Fig. 5e**). Interestingly, although the predicted contribution of EICE motifs was higher in the preGC-2 state, they were also predicted to contribute in the PB state albeit with an altered nucleotide contribution score profile with less emphasis on the ETS-half of the composite element. This was consistent with the increased activity of PRDM1 which is up-regulated in PBs and known to compete with PU.1/SPIB for binding at EICE motifs in murine B cells^42^. Thus, in strong agreement with the MIRA-CellOracle GRN modeling (**Fig. 4e**), the ChromBPNet analyses support a model in which the counterbalance between IRF4/IRF8 and their binding partners at AICE, EICE and ISRE motifs plays a critical role in determining human B cell fate choice.

### Single-cell perturbation analysis of counteracting PB and preGC TFs

The ChromBPNet analyses (**Fig. 5e**), combined with the simulated and experimental TF perturbation screens (**Fig. 4e**), strongly suggested that preGC versus PB fate specification depends on distinct cooperative and antagonistic interactions among IRF4, IRF8, BATF, SPIB and BLIMP1 via their binding at AICE, EICE and ISRE motifs. To test this hypothesis at genomic resolution, we analyzed the consequences of knocking out *IRF4*, *SPIB*, *BATF* or *PRDM1* (BLIMP1) in activated B cells by scRNA-seq. The experimental design involved performing CRISPR RNP nucleofections at D2 followed by scRNA-seq at D4 and D6 as the cells diverge into PB or preGC trajectories (**Fig. 6a**, *top left*). Following MIRA gene expression topic modeling and Leiden clustering, we identified three major clusters corresponding to ActB, preGC and PB states (**Fig. 6a**, *top middle*). While the ActB cluster was exclusively derived from the Day 4 timepoint, the preGC and PB clusters were primarily composed of Day 6 cells and expressed high levels of their respective marker genes (*LMO2 and AICDA* for preGCs and *XBP1* and *JCHAIN* for PBs) (**Fig. 6a** and **Extended Data Fig. 6a**). In agreement with the immunophenotyping analysis (**Fig. 4e**), BATF- and SPIB-KOs displayed moderate skewing towards PB differentiation (**Fig. 6b**). In contrast, PRDM1- and IRF4-KO cells were severely impaired for PB formation and biased towards the preGC fate. Notably, whereas PRDM1-KO cells generated preGCs that were indistinguishable from their control counterparts, IRF4-KO cells were nearly completely segregated from other cells in the preGC cluster, likely representing a stalled preGC transition state (**Fig. 6a**, *bottom*). This was consistent with the established dual role of IRF4 in controlling both PB and GCBC differentiation^5,6^.

**Fig. 6.**
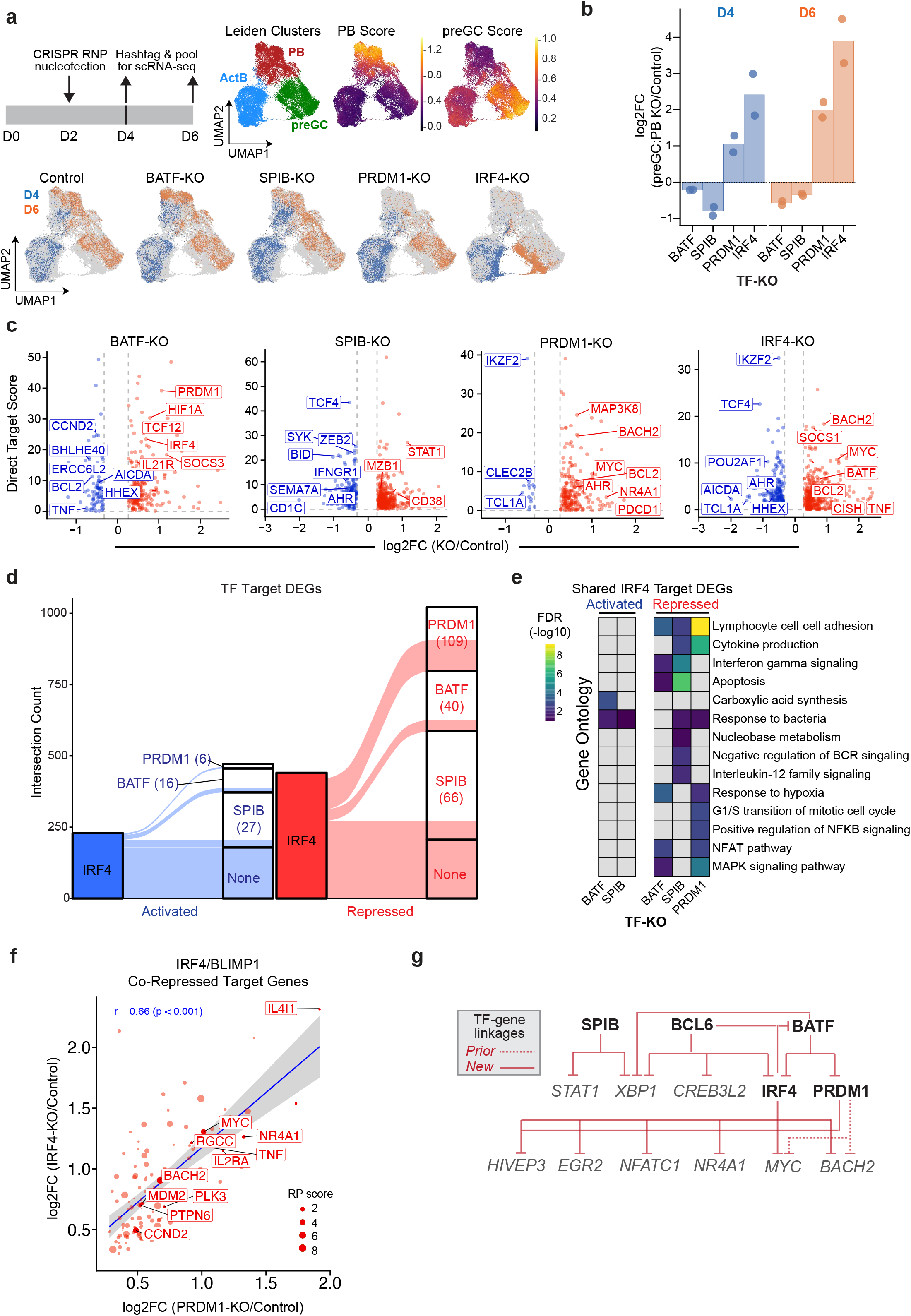
Single-cell perturbation analysis of counteracting PB and preGC TFs. **(a)** Experimental strategy for profiling TF perturbations by scRNA-seq (*top left*). Naïve B cells (Donor 1) were nucleofected with CRISPR RNPs at Day 2 and harvested at Days 4 and 6 for hash - tagging and pooled scRNA-seq. Aggregated UMAP showing Leiden clusters and PB/preGC scores (*top right*). UMAPs showing cells from each TF-KO condition colored by timepoint (*bottom*). **(b)** Plot displaying log2 fold change in preGC:PB ratio at Day 6 for each TF-KO condition versus control (n = 2 independent experiments using Donor 1 cells). **(c)** Volcano plots showing log2 fold change versus direct target score for concordant DEGs (|FC| > 1.2, FDR < 0.05 in both replicates) in each TF-KO condition. Direct target score = sum of MIRA RP values for ATAC-seq peaks containing ChromBPNet seqlets matching the perturbed TF’s binding motif. Dotted lines indicate the |FC| = 1.2 threshold. **(d)** Intersection of IRF4-KO DEGs with BATF-, SPIB-, and PRDM1-KO DEGs. Numbers indicate concordantly co-regulated genes. Activated (blue) and repressed (red) genes are shown separately. **(e)** Gene ontology enrichment analysis of IRF4 co- regulated DEG modules. Heatmap shows −log10(FDR) for pathways enriched among genes co- activated or co-repressed by IRF4 and indicated partner TFs. (**f**) Scatter plot displaying IRF4 and PRDM1 co-repressed genes linked to at least one EICE or ISRE motif seqlet (n = 109 genes). Sizes of the circles indicate the RP score for the EICE or ISRE seqlet-containing peaks at indicated target genes. (**g**) Schematic summarizing representative confirmatory and novel TF-TF cross-regulatory linkages collectively recovered from the TF perturbation scRNA-seq experiments.

To uncover gene expression programs that are cooperatively or antagonistically regulated by these TFs, we analyzed the differentially expressed genes (DEGs) within the Day 4 ActB cluster. By focusing our analysis on bipotent ActB cells, we increased the likelihood of detecting TF-controlled gene modules that initiate programming of preGC or PB fates. Profiling of the perturbed transcriptomes 48 hours post-RNP nucleofection also minimized secondary effects, thereby enriching for direct TF target genes. We restricted our analysis to DEGs that were concordantly significantly up- or down-regulated (|FC| > 1.2, FDR < 0.05) between the two replicates (Pearson correlation coefficient, r = 0.88 to 0.97) (**Extended Data Fig. 6b**). IRF4-KO cells displayed the most DEGs (n=720), followed by SPIB-KO (n=594), BATF-KO (n=297), and PRDM1-KO (n=217).

To identify genes that were likely to be directly regulated by each of the TFs (i.e., targets within their cognate sets of DEGs), we integrated the MIRA RP models with those from ChromBPNet. The former predicted cCREs for each DEG based on the single-cell correlations between gene expression levels and accessibility of the cCREs (**Fig. 3a**), whereas the latter inferred sites of TF binding and action within those cCREs based on predicted base-pair contributions to chromatin accessibility profiles (**Fig. 5a**). To infer which cCREs are regulated by IRF4 or one of its partner TFs in activated B cells, we used the Day 3 ActB ChromBPNet model to identify cCREs that contain seqlets whose contribution weight matrices were matched with relevant annotated TF motifs (see Methods). Unlike simple motif scanning, this approach discerned which TF motif instances likely contribute to the accessibility of a given cCRE in a particular cell state. We identified all seqlet instances within the union set of ATAC-seq peaks for AP-1 (BATF), ETS (SPIB), ISRE (IRF4), PRDM1 (BLIMP1), AICE (BATF::IRF4), and EICE (SPIB::IRF4, BLIMP1) (**Extended Data Fig. 6c**). These motif seqlets were then intersected with cCREs linked to DEGs via the RP models. This enabled quantitation of the number of seqlet- cCREs and their regulatory potential, resulting in a direct target score for each TF-gene pair (sum of individual seqlet-containing cCRE RP values within a given RP model) (**Fig. 6c**). For the delineation of direct target genes of a given TF, we identified DEGs of that TF, each of which contained at least one cognate ChromBPNet seqlet within a RP model-linked cCRE. This resulted in delineation of SPIB- (n=562), BATF- (n=289), IRF4- (n=629), and PRDM1- (n=201) regulated genes that were putative direct targets (**Supplementary Table 12**). To evaluate the quality of the TF-target gene linkages in the MIRA assembled base-GRN model (**Fig. 3**), we tested for enrichment of TF KO DEGs in corresponding TF-target gene sets (**Extended Data Fig. 6d**). This revealed that all four TF-KO DEGs were enriched in the MIRA-predicted base-GRN TF-gene linkages (BATF: 67 validated predicted target genes, odds ratio = 3.4, p = 3.8e-12; SPIB: 69 validated predicted target genes, odds ratio = 2.5, p = 4.17e-9; PRDM1: 10 validated predicted target genes, odds ratio = 1.68, p = 0.09; IRF4: 136 validated predicted target genes, odds ratio = 3.8, p = 1.89e-26). Notably, MIRA-based TF-gene predictions derived from ChIP-seq peak ISD modeling substantially outperformed the default Cicero-based approach using co-accessibility and TF motif enrichment.

Based on our hypothesis that IRF4 co-regulates divergent B cell programs through its combinatorial activity with distinct partner TFs, we intersected IRF4 target genes with those of its putative binding or cooperating partners. Using enrichment analyses (hypergeometric tests), we identified significant overlaps between the IRF4 activated or repressed target genes and those of its putative partners (adjusted p = 1.5e5 and 2.5e-96 for PRDM1 co-repressed and -activated genes; adjusted p = 6.6e-10 and 2.6e-16 for BATF co-repressed and -activated genes; adjusted p = 2.5e-13 and 4.2e-25 for SPIB co-repressed and -activated genes) (**Fig. 6d**, **Supplementary Table 13**). Gene ontology analysis revealed enrichment in immune-related pathways for all IRF4 co-regulated genes except for IRF4/PRDM1 co-activated genes which had very small sample sizes (**Fig. 6e**, **Supplementary Table 14**). IRF4 and BATF co-activated genes were enriched for pathways associated with carboxylic acid synthesis, whereas both BATF and SPIB co-activated genes with IRF4 that were enriched for response to bacteria. BATF and SPIB both specifically co-repressed interferon gamma signaling and apoptosis pathways with IRF4, while SPIB-IRF4 co-repressed genes were uniquely enriched for IL12 signaling and BCR signaling pathways. In contrast, IRF4 and PRDM1 (BLIMP1) co-repressed genes were uniquely enriched for positive regulators of NFκB signaling (*CCR7*, *TNF*, *TNFRSF10B*, *NAMPT*), G1/S transition (*MYC, CCND2, PLK3, RGCC*), and lymphocyte cell-cell adhesion (*CD80, CD70, SLAMF1, CD274, SEMA7A*).

In activated murine B cells that are differentiating into PBs, IRF4 and BLIMP1 have remarkably similar binding profiles with over half of all BLIMP1-bound regions co-bound by IRF4^42^. These co-bound regions are highly enriched for EICE motifs (GGAA-NN-GAAA) and ISRE-like motifs (GAAAG-N-GAAAG)^45^, raising the possibility that IRF4 and BLIMP1 cooperatively repress shared target genes by binding the same cCREs. To investigate whether IRF4 and BLIMP1 function as co-repressors in activated B cells, we compared the changes in gene expression in IRF4-KO and PRDM1-KO cells. Concordantly up-regulated genes that were linked to EICE or ISRE seqlets had a significant positive correlation (Pearson’s correlation, r = 0.66, p = 2.8e-15, n = 109 genes) (**Fig. 6f**), consistent with cooperative or convergent repression via shared cCREs. Among these co-repressed genes were known regulators of cell cycle and proliferation (MYC, RGCC, MDM2, PLK3, PTPN6, IL2RA) as well as regulators of B cell activation and differentiation (BACH2, EGR2, NR4A1, TNF, IL4I1). Gene set enrichment analysis confirmed that MYC target genes were significantly enriched among genes up-regulated in both IRF4-KO (NES = 2.23, FDR ∼ 0) and PRDM1-KO (NES = 2.28, FDR ∼ 0) cells (**Extended Data Fig. 6e,f**). Together with the enrichment of G1/S transition genes among IRF4/BLIMP1 co-repressed targets (**Fig. 6e**), these results suggest that IRF4 and BLIMP1 concertedly restrain cell cycle progression while also silencing activated B cell programs.

Additionally, the CRISPR perturbation experiments collectively revealed extensive evidence of previously undocumented TF-TF cross-regulation (**Supplementary Table 15**). These include negative regulation of PB-associated TFs (*STAT1*, *XBP1*, *CREB3L2, IRF4*) by SPIB and/or BCL6 and activated B cell and GCBC-associated TFs (*HIVEP3*, *EGR2*, *NFATC1*, *NR4A1*, *MYC*, *BACH2*) by IRF4 and BLIMP1 (**Fig. 6g**). Interestingly, BATF was repressed by IRF4 and PRDM1, while IRF4 reciprocally repressed BATF, providing evidence for a previously undocumented negative feedback loop that operates alongside previously established ones to generate robust bistability^8^. Together, these results (1) provide rigorous fine-grained testing of the TF-gene linkages underlying the B cell GRN models, (2) validate hundreds of predicted target genes for four core TFs and (3) provide new insights into how these TFs combinatorially promote divergent B cell fates.

### Predicting and interpreting B cell state-specific effects of autoimmune-associated variants

B cells are central to the pathogenesis of a wide range of autoimmune diseases^3^. Variants associated with autoimmune diseases are overwhelmingly located in non-coding genomic regions, making it difficult to predict and interpret their potential contribution to disease^46^. Further complicating matters, it remains difficult to determine which cell-type and which genes a particular variant may impact. To address these challenges and nominate functional autoimmune disease variants and their impact on gene expression programs in human B cells, we leveraged our ChromBPNet single base-pair chromatin modeling to predict and interpret B cell state-specific variant effects (**Fig. 7a**).

**Fig. 7.**
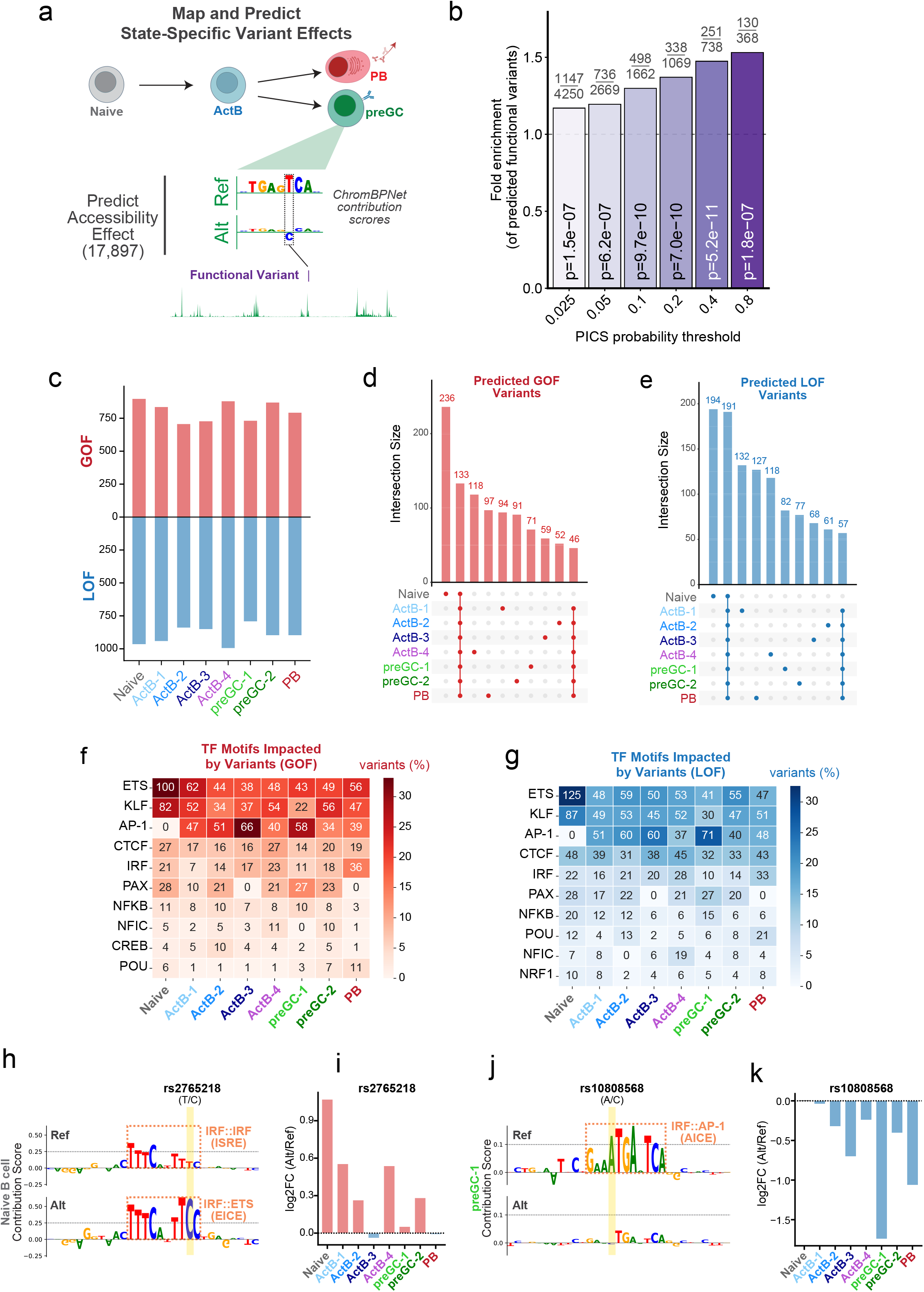
Predicting and interpreting B cell state-specific effects of autoimmune-associated variants. **(a)** Workflow for predicting B cell state-specific variant effects. Fine-mapped autoimmune-associated variants from the PICS2 GWAS catalogue that mapped to B cell OCRs (n = 17,897) were analyzed with state-specific ChromBPNet models to predict variant effects on chromatin accessibility. (**b**) Enrichment of ChromBPNet-predicted functional variants among fine-mapped PICS2 autoimmune variants. Fisher’s exact test was performed at varying PICS probability thresholds, using all OCR-residing variants with a PICS probability < 0.025 and all ChromBPNet variants with p > 0.05 as the background sets. Numbers above the plot indicate the number of variants that satisfy both the PICS2 and ChromBPNet p-value thresholds over the total number of variants at each threshold. The minimum ChromBPNet p-value across all 8 B cell states was used in all calculations. (**c**) Number of ChromBPNet-predicted GOF and LOF variants per B cell state. (**d,e**) Upset plots displaying the number of shared and state-specific ChromBPNet-predicted GOF (**d**) or LOF (**e**) variants. The top 10 most numerous intersection sets are displayed. (**f**,**g**) Heatmaps showing the number of predicted GOF (**f**) or LOF (**g**) variants impacting each TF motif family across B cell states. Cell values display variant counts and the color scale indicates the percentage of all variant-motif pairs within each state. CWMs for alternative (GOF) or reference (LOF) alleles were matched to known TF motifs using Tomtom. Top 10 most frequently impacted motif families are shown per state. **(h-k)** Examples of state-specific variant effects. **(h)** Contribution score logos for rs2765218 reference (*top*) and alternative (*bottom*) alleles in naïve B cells, showing conversion of an ISRE to an EICE motif. **(i)** Predicted log2 fold change in local accessibility (2 kb window) for rs2765218 across B cell states. **(j)** Contribution score logos for rs10808568 showing disruption of an AICE motif in the preGC-1 state. **(k)** Predicted accessibility effects for rs10808568 across B cell states.

We first mapped 17,897 of the 122,653 unique single nucleotide variants associated with one or more autoimmune diseases from the PICS2-curated GWAS catalogue^47^ to the union set of open chromatin regions from our single-cell multiome dataset (**Fig. 2a**). We then applied our B cell state-specific ChromBPNet models (**Fig. 5**) to predict the local chromatin accessibility effects (∼2kb windows) of each variant in each of the eight major B cell states (**Supplementary Table 16**). This identified 4,248 variants predicted to significantly impact (p < 0.05) local chromatin accessibility in at least one B cell state. To determine if these predicted functional variants were associated with increased autoimmune disease risk, we compared the ChromBPNet variant predictions to fine-mapping results from PICS2, which estimates the probability that an individual variant is causal given the haplotype structure and locus association patterns. Predicted-functional variants (p < 0.05) were significantly enriched among fine-mapped autoimmune variants at increasing PICS probability thresholds, reaching statistical significance above a PICS probability of 0.025 and ∼1.5-fold enrichment at a PICS probability > 0.8 (p = 1.8e-7) (**Fig. 7b**).

To investigate whether our B cell ChromBPNet models accurately predict the effect size and directionality of variant effects on chromatin accessibility in B cells, we compared the predicted changes in chromatin accessibility to the observed allelic skewing reported in caQTL analysis of primary B cells^48^ and EBV-immortalized B cell lymphoblast cell lines (LCLs)^49^. Predicted accessibility effects were strongly correlated with observed allelic effect sizes and directionality in primary B cell caQTL analysis (Pearson correlation coefficients: 0.42-0.63; Spearman correlation coefficients: 0.34-0.6) (**Extended Data Fig. 7a**). Similarly, the predicted directionality was highly concordant with allelic skew observed for LCL caQTL analysis (**Extended Data Fig. 7b**). This demonstrates that the B cell ChromBPNet models reliably predict the effects of variants on chromatin accessibility in B cells.

To determine if ChromBPNet-predicted functional variants were also associated with effects on transcriptional activity, we compared our ChromBPNet predictions to expression-modulating variant (emVar) calls from six LCL MPRA datasets^50–52^, including a new dataset generated for this study (**Supplementary Tables 17 and 18**). Predicted-functional variants were enriched for emVars across the six LCL MPRA datasets, reaching ∼2.2-fold enrichment at p < 0.001, including 13 ChromBPNet-predicted functional emVars unique to the new MPRA dataset generated for this study (**Extended Data Fig. 7c,d**). This new dataset comprised 54 autoimmune-associated variants residing in B cell OCRs mapped in this study, of which 14 significantly altered transcriptional activity in GM12878 cells (FDR < 0.1) (**Supplementary Table 18**).

Each B cell state harbored approximately 700 to 1,000 predicted gain-of-function (GOF) and loss-of-function (LOF) variants (**Fig. 7c**). Comparison of B cell state-specific predictions revealed that only 133 GOF and 191 LOF variants were shared across all eight B cell states (**Fig. 7d,e).** Notably, 52 to 236 GOF (818 total) and 61 to 194 LOF (859 total) variants were predicted to have significant effects exclusively in a single B cell state. We next investigated which ChromBPNet-predicted functional variants were likely to specifically affect only B cells rather than other autoimmune-relevant immune cell types. Using published ChromBPNet models trained on ATAC-seq data from activated CD4 T cells, CD8 T cells, and activated dendritic cells, we found that 48% of GOF (997) and 47% of LOF (1,028) variants were predicted to affect chromatin accessibility in at least one B cell state but in none of these other activated immune cell-types (**Extended Data Fig. 7e,f**; **Supplementary Table 16**). These results suggest that the impact of many autoimmune disease variants is highly cell-type and cell-state dependent.

Given that TF motif and regulatory activity vary substantially across B cell states (see **Fig. 5d**), we hypothesized that these state-specific variant effects were driven by differential TF activity. To test this, we used ChromBPNet contribution score weight matrices (CWMs) for either alternative or reference allele sequences to identify the accessibility-contributing TF motifs that would be created or disrupted by GOF or LOF variants, respectively. Strikingly, GOF and LOF variants impacted dramatically different TF motif families across B cell states (**Fig. 7f,g** and **Supplementary Table 19**). While predicted functional variants at CTCF motifs were uniformly represented across all B cell states, variants at ETS, KLF and AP-1 related motifs exhibited divergent activity in naïve versus activated/differentiated B cell states. Specifically, while variants in naïve B cells predominantly affected ETS and KLF related motifs, the frequency of variants impacting AP-1 motifs dramatically increased in activated/differentiated B cell states. Notably, the variant-impacted motif family frequencies also differed between preGC and PB states. While the AP-1 and PAX motifs composed a large fraction of variant-impacted motifs in the preGC-1 state, variants in the PB state completely lacked PAX related motifs but were more enriched for IRF-related motifs, which include IRF4 and PRDM1 binding sites, and POU-related motifs, which include Oct-1,2 binding sites. To further resolve the state-specific nature of variant impacts on TF motifs, we repeated this analysis for GOF or LOF variants that were specific to a single-cell state. Strikingly, while variants that were shared across all states were predicted to impact a wide range of TF motifs, variants that were specific to naïve B cells predominantly impacted ETS and KLF motifs. Furthermore, variants that were predicted to impact all B cell states, with the exception of naïve B cells, were predominantly predicted to impact AP-1 motifs, highlighting a partitioning of autoimmune variant impact on ETS versus AP-1 motifs in naïve versus activated/differentiated B cell states.

To illustrate state-specific variant effects, we highlight two examples (**Fig. 7h-k**). The first example, rs2765218, converted an ISRE motif into an EICE motif (**Fig. 7h**), with predicted impact restricted to naïve and early activated states (**Fig. 7i**), consistent with the transition from ETS- dominated to IRF-dominated regulatory activity during differentiation. The second example, rs10808568, disrupted an AP-1::IRF composite element (AICE) motif (**Fig. 7j**) which was predicted to mostly impact the preGC-1 state (**Fig. 7k**), consistent with heightened cooperative activity of AP-1 and IRF family TFs during B cell activation and GC initiation.

### Predicted functional variants are enriched for eQTL variants linked to autoimmune disease programs

We next investigated which ChromBPNet-predicted functional variants were also cis-acting expression QTL (cis-eQTL) variants that were fine-mapped across six different LCL datasets (**Extended Data Fig. 8a**). Of the 3,698 cis-eQTL variants that resided in B cell OCRs and were associated with one or more autoimmune diseases, 783 were predicted to have a significant impact on chromatin accessibility (**Fig. 8a**). These ChromBPNet-predicted functional variants were significantly enriched among fine-mapped cis-eQTL variants at increasing posterior inclusion probability (PIP) thresholds (**Fig. 8b**), reaching ∼2-fold enrichment at PIP > 0.8. Additionally, of the 783 ChromBPNet-predicted functional eQTL variants, 210 carried at least one additional line of functional support (LCL caQTL and/or MPRA emVar), with 21 of those meeting both criteria (**Fig. 8c**). Notably, around 20% of the 783 predicted functional eQTL variants also had a PICS probability > 0.025, indicative of candidate causal disease risk variants (**Extended Data Fig. 8b**).

**Fig. 8.**
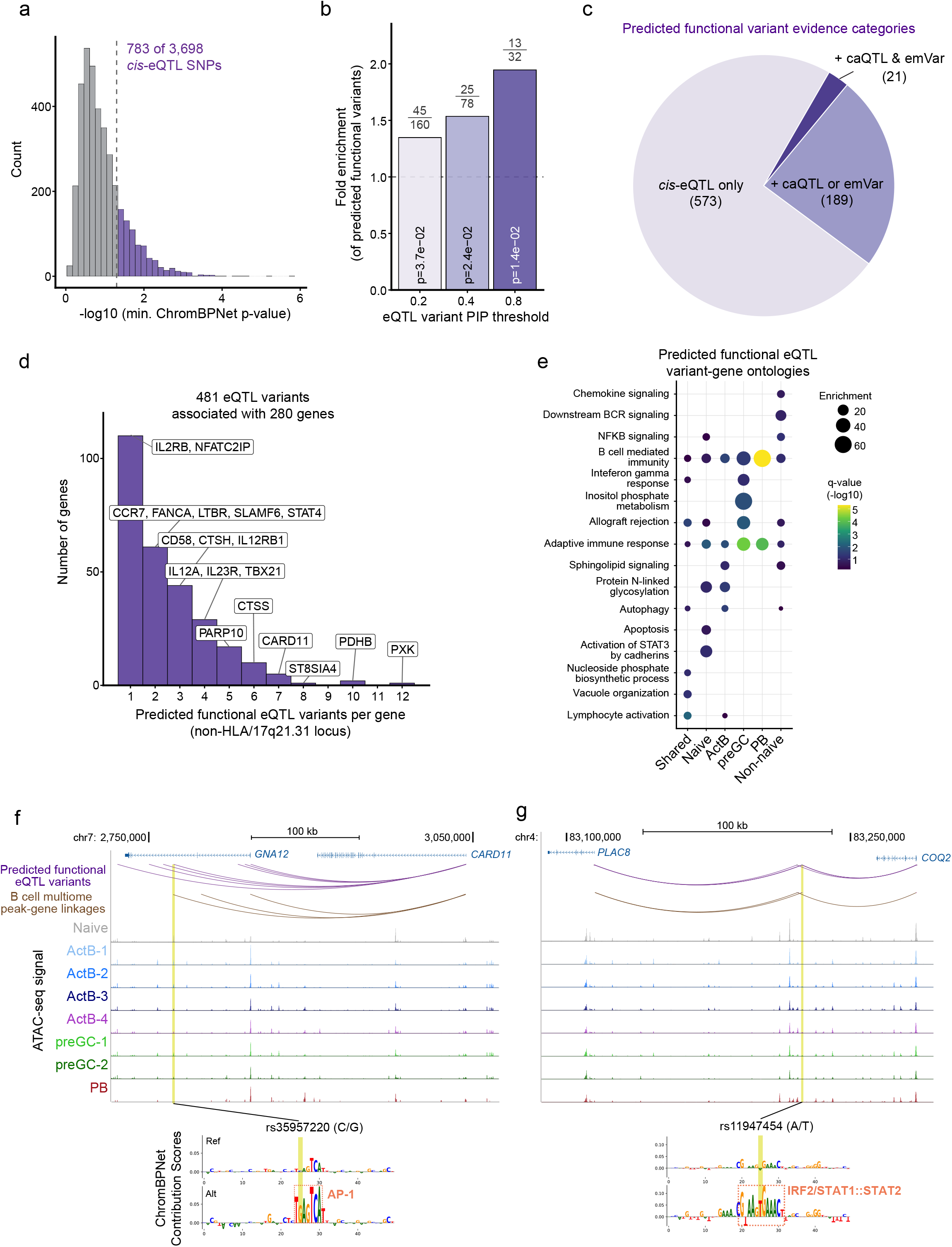
Predicted functional variants are enriched for eQTL variants linked to autoimmune disease programs. (**a**) Distribution of minimum ChromBPNet p-values for 3,698 *cis*-eQTL variants mapped to B cell OCRs. eQTL variants with a minimum ChromBPNet p < 0.05 (dotted vertical line) were considered as predicted to be functional by ChromBPNet for downstream analysis. (**b**) Enrichment of ChromBPNet-predicted functional variants among LCL cis-eQTL credible variants. Fisher’s exact test was performed at varying eQTL posterior inclusion probability (PIP) thresholds, using all OCR-residing eQTL variants with PIP < 0.2 and all ChromBPNet variants with p > 0.05 as the background sets. Numbers above the plot indicate the number of variants that satisfy both the eQTL PIP and ChromBPNet p-value thresholds over the total number of tested variants at each eQTL PIP threshold. The minimum ChromBPNet p-value across all 8 B cell states was used in all calculations. (**c**) Pie chart displaying the number of ChromBPNet- predicted functional eQTLs that have additional levels of evidence (caQTL or emVar from LCL studies; see Extended Data Fig. 7). (**d**) Histogram displaying the distribution of predicted functional eQTL variants per gene. Variants residing in loci with frequent structural inversions/duplications (HLA & 17q21.31) were excluded from the analysis due to fine-mapping limitations in these regions. Select genes that are associated with autoimmune diseases/traits are annotated. (**e**) Dot plot displaying enrichment (size) and FDR (color) for select gene ontologies that are associated with autoimmune disease traits. Each column represents sets of genes that are associated with eQTL variants that were predicted to be functional in all B cell states (“Shared”) or specific to one or more indicated B cell states. (**f**,**g**) Representative predicted functional eQTL variants that are also LCL caQTLs and reside in peaks that are linked to corresponding eQTL genes. (**f**) Genome browser view of *CARD11* locus with inset displaying ChromBPNet predictions at rs35957220. The gain-of-function AP-1 motif is annotated with a dotted box. (**g**) Genome browser view of the *PLAC8*/*COQ2* locus with inset displaying ChromBPNet predictions at rs11947454. The gain-of-function IRF2/STAT1::STAT2 motif is annotated with a dotted box.

We next investigated the genes whose expression was associated with the predicted functional eQTL variants. The 783 predicted functional eQTL variants were associated with 332 unique genes, with each gene being linked to up to 78 unique eQTL variants (**Extended Data Fig. 8c**). The outlier genes associated with the most eQTL variants all resided in either the HLA or 17q21.31 loci, which dominated the eQTL variants found on chromosomes 6 and 17, respectively (**Extended Data Fig. 8d**). After excluding variants found in these highly complex regions that contain frequent inversions and duplications, which can complicate interpretation of fine-mapping results, there were 481 eQTL variants associated with 280 unique genes (**Fig. 8d** and **Supplementary Table 20**). The ChromBPNet-predicted functional eQTL variants were linked to genes that have documented functional roles in autoimmune diseases (*STAT4*, *IL12A*, *IL23R*, *IL12RB1*, *IL2RB*, *CD58*, *CTSS*, *TBX21*, *PXK*, *CARD11*). Notably, most genes were associated with only 1-3 ChromBPNet-predicted functional variants, despite eQTL credible sets often containing many candidate causal variants. This illustrates how integrating fine-mapping with sequence-to-function modeling can narrow multi-variant credible sets to a small number of prioritized causal candidates.

The genes linked to the predicted functional eQTL variants were associated with wide ranging autoimmune diseases (**Extended Data Fig. 8e**) and were enriched for many gene ontology pathways, which have previously been linked to autoimmune disease traits (e.g. chemokine signaling, downstream BCR signaling, NFKB signaling, interferon gamma responses, autophagy, vacuole organization) (**Fig. 8e** and **Supplementary Table 21**). For example, there were 7 predicted functional eQTL variants associated with the expression of *CARD11*, a critical signaling scaffold protein that modulates downstream B cell receptor pathways (**Fig. 8f**). One of the eQTL variants (rs35957220) resided in an OCR whose accessibility was significantly correlated with *CARD11* expression in our B cell multiome dataset (**Supplementary Table 21)**. Our B cell ChromBPNet models predicted that the eQTL variant would create a functional AP-1 motif that would increase the OCR’s accessibility in activated B cells. A second example can be observed at the *PLAC8*/*COQ2* locus, where three eQTL variants reside in OCRs whose accessibility was significantly correlated with the expression of either *PLAC8* (a key regulator of autophagy) or *COQ2* (an enzyme required for coenzyme Q10 production) (**Fig. 8g** and **Supplementary Table 21**). Our B cell ChromBPNet model predicted that one eQTL variant in particular (rs11947454) would create a functional IRF2 or STAT1::STAT2 motif that would increase the OCR’s accessibility in activated B cells. Altogether, these results demonstrate that the ChromBPNet variant effect predictions are functionally relevant and highlight the power of using such high-resolution predictions to interpret and prioritize novel variants/genes for characterization studies.

## Discussion

This work provides multi-scale, state-specific GRNs governing the bifurcation of human B cells into PB and preGC trajectories, at a scale and resolution not previously available. We demonstrate the utility of this resource at two levels: First, we apply a top-down approach for nominating novel TFs, CREs and regulatory linkages that control naive B cell activation and differentiation into PB or preGC trajectories. Second, we apply a bottom-up approach for predicting and interpreting the impact of non-coding autoimmune variants on B cell state-specific chromatin accessibility and gene expression. Importantly, the GRN models were rigorously tested at multiple scales including TF-to-fate and TF-to-gene predictions. This testing demonstrated that our integration of orthogonal computational methods (MIRA regulatory potential modeling coupled with CellOracle GRN inference) significantly outperforms standard approaches based on Cicero-derived base-GRNs. Complete state-specific GRN models and simulations for 124 central TFs are available through our interactive web application (https://pitt-csi.shinyapps.io/humanBcellGRN/), nominating previously unrecognized regulators including pro-GC TFs (BCL11A, HIVEP3, TFEC, IKZF2, PBX3) and pro-PB TFs (RUNX2, CREB3L2, BACH1, KLF2). Our integration of base-pair resolution ChromBPNet modeling with peak-to-gene analysis enables systematic predictions of how non-coding variants associated with autoimmune diseases may impact transcription factor binding and gene regulation across diverse B cell states. Thus, the assembled GRNs composed of multi-scale predictions and an accompanying interactive web application provide a foundational resource for analyzing human B cell states in health and autoimmune diseases^38,51^.

A limitation of GRN modeling arises when inferred networks are evaluated only within the system that generated the training data. To test whether our framework generalized beyond the *in vitro* human B cell system, we applied the identical MIRA and CellOracle pipeline to multiome data from the Human Tonsil Atlas and assembled GRNs for the corresponding *in vivo* B cell states. Chromatin accessibility landscapes across the datasets substantially overlapped in a state-specific manner. TF centrality scores and TF-gene edge weights were significantly correlated between corresponding *in vitro* and tonsil states, and in silico perturbations recovered the same regulators in both settings, including established drivers (PB/PC: IRF4, PRDM1; preGC/GCBC: BCL6, IRF8, MEF2C, PAX5, SPIB, BATF) as well as candidate regulators newly nominated herein (PB/PC: RUNX2, CREB3L2, KLF2; preGC/GCBC: MEF2A, TFEC, IKZF1/2, PBX3, HIVEP3). While the global correlation of perturbation scores was modest in magnitude, the concordance was driven by directionally consistent recovery of these key regulators. Predicted BCL6 target genes were further corroborated experimentally, with BCL6-KO up- regulated genes significantly enriched among genes predicted to be repressed by BCL6 in both the *in vitro* preGC and tonsil GCBC models. Thus, the computational genomics human B cell resource encompasses both temporally and developmentally resolved *in vitro* and *in vivo* tonsil states.

The expansion of single-cell analyses that simultaneously profile transcriptional and chromatin landscapes has propelled the widespread application of GRN modeling across diverse developmental and cellular contexts^53^. However, GRN modeling has been limited by several issues addressed herein. First, instead of relying solely on co-accessibility patterns to connect candidate CREs (cCREs) to target gene promoters, we apply MIRA RP modeling which infers the regulatory potential of cCREs neighboring a gene based on co-variation of cCRE accessibility and gene expression. Second, rather than predicting TF binding and action on cCREs simply based on TF motif enrichment and/or occurrence, which cannot distinguish different structurally related TFs, we utilize the CistromeDB database as a source of over six thousand uniformly processed ChIP-seq datasets. MIRA ISD modeling is then used to identify likely TF-gene connections based on simulating the removal of cCREs with evidence of TF binding in related cellular contexts. Third, rather than assigning the same TF-gene linkages to all cell states of a given system, we applied CellOracle GRN modeling to transform cell-state agnostic and qualitative linkages into state-specific and quantitative TF-gene linkages. Collectively, integration of these three modeling approaches allows us to simultaneously anchor the TF-gene linkages to physical TF-specific binding events and to infer the strength and directionality of linkage reconfigurations during a dynamic differentiation process. Importantly, the improved performance of GRN models based on an integration of MIRA with CellOracle was demonstrated with two types of experiments that tested the impact of TF perturbations on cell fate determination as well as on their target genes (see below).

The assembled GRNs provide comprehensive predictions of TFs and their target genes that drive human B cell responses and account for their functions. Previously, such in silico perturbations have not been systematically tested with corresponding experimental perturbations. Here, we tested the impact of perturbing 31 TFs that were predicted to differentially impact the generation of preGCs and PBs. These results represent, to our knowledge, the first attempt to systematically and quantitatively test a large panel of TFs in controlling human B cell differentiation into preGC and PB trajectories, which builds upon the PB versus GC models previously established in murine systems and other *in vitro* human B cell systems^5,10,11^. The integrated computational framework (MIRA and CellOracle) demonstrated excellent concordance between in silico perturbations and their experimental analyses demonstrating the quality of the assembled GRNs. Notably, CellOracle and other *in silico* perturbation approaches have performed less well in benchmarking analyses^54^. This has primarily been attributed to limitations of modeling assumptions for in silico perturbations. However, our analyses suggested that higher-quality GRN assemblies can significantly mitigate issues with corresponding modeling assumptions. In addition to demonstrating the effectiveness of our GRN model predictions, the results from the predict-perturb-test framework^55^ both corroborate and substantially extend our previous knowledge of murine B cell GRNs^7,56,57^.

Our systematic in silico and *in vitro* perturbations provide converging computational and experimental support that IRF4 and PRDM1 are the dominant regulators of PB differentiation in the human system, consistent with perturbation screens performed in murine systems^58–61^. In contrast, the preGC fate is controlled by many different TFs, each with relatively moderate effects on preGC generation. This is likely in part due to the limited phenotypic space which can be assayed *in vitro*. Of note, while BCL6 was predicted to have a strong positive effect on the preGC:PB ratio, the BCL6-KO cells exhibited only a mild effect on the generation of this *in vitro* preGC state based on low-dimensional phenotypic marker-based analysis. However, scRNA-seq analysis of BCL6-deficient cells revealed profound disruption of the preGC transcriptional program, which failed to upregulate genes distinguishing tonsil GCBCs from activated B cells. This demonstrates that BCL6 is essential for establishing the preGC transcriptional state. Importantly, our *in vitro* system generates GCBC precursor cells (preGCs) rather than mature GCBCs. The preGC state is transcriptionally intermediate between activated B cells and tonsil GCBCs, as demonstrated by Harmony integration analysis showing these cells bridge tonsil ActB and GCBC populations. Generation of mature GCBCs would likely require reconstitution of the complex cellular interactions provided by the germinal center microenvironment, including sustained interactions with T follicular helper cells and follicular dendritic cells. Consequently, while our data establish that the *in vitro* preGC transcriptional state is BCL6-dependent, they cannot establish whether this preGC state has the capacity to generate bona fide GCBCs. Addressing that question would require systems capable of supporting complete GC reactions with affinity maturation and positive selection. Nevertheless, the preGC state we describe is BCL6-dependent and recapitulates the transcriptional and chromatin features of GCBC differentiation, providing a tractable system for investigating the molecular mechanisms that initiate the GCBC trajectory and programming. The absence of single TF perturbations in our experimental system that dramatically decrease the human preGC:PB ratio by surface markers mirrors results of CRISPR screens in murine B cells^58–60^, suggesting that preGC/GCBC formation is likely regulated by multiple, partially redundant factors that converge on a BCL6-dependent transcriptional program.

Here, by systematic profiling of perturbations of IRF4 and its potential binding partners, we delineated gene programs that are co-regulated by various combinatorial pairs in activated B cells that could give rise to PBs or preGCs. The most striking co-regulatory relationship was between IRF4 and PRDM1, which we found initially function to co-repress genes associated with B cell activation, survival and proliferation programs. This contrasts with the cell migration and antibody secretion gene programs that IRF4 and PRDM1 promote during PB and plasma cell differentiation^6,42^. The results strongly suggest that the specification of the PB fate is initiated by repression of gene programs that are needed to sustain the preGC trajectory. Notably, while IRF4 and PRDM1 appear to reinforce each other’s expression in the PB state^42^, we did not find any experimental evidence for such reciprocal positive feedback in the ActB state. Instead, the GRN model and its testing revealed that *IRF4* and *PRDM1* are repressed by a common up-stream regulator, BATF, that promotes the GCBC fate. Thus, diminished BATF expression results in simultaneous IRF4 and PRDM1 upregulation, which feed back to repress their own repressors, BATF and BACH2. We have previously demonstrated that such a reciprocal negative feedback structure can generate robust bistability^8^. Future studies will be required to comprehensively elucidate the additional layers of regulation that tune this bistable switch to a IRF4^hi^BLIMP1^hi^ regulatory state.

Our ChromBPNet-based analysis of autoimmune disease-associated variants revealed a striking partitioning of genetic risk across naïve, activated and differentiated regulatory compartments. Variants predicted to impact ETS and KLF family motifs showed substantial effects in naïve B cells, consistent with their occurrence in constitutively accessible regulatory elements. In stark contrast, variants affecting AP-1 family motifs were absent from naïve B cells but emerged dramatically upon B cell activation, reaching maximal impact in ActB and preGC states. This pattern suggests that AP-1-associated autoimmune variants reside within activation-induced open chromatin regions that are initially inaccessible and thus functionally silent until antigen receptor and co-stimulatory signals trigger chromatin remodeling. These findings extend prior observations in T cells and macrophages demonstrating enrichment of autoimmune variants in stimulation-responsive CREs^62–64^. Notably, Alasoo et al. identified “enhancer priming” as a mechanism whereby variants alter chromatin accessibility in naïve macrophages, but the effects of such variants on gene expression are observed upon stimulation, a pattern consistent with our variant effect predictions at ETS/KLF motifs in naïve B cells. However, our AP-1-associated variants represent a distinct mechanistic class: de novo activation-induced elements where the chromatin itself must first become accessible before the variant can exert any regulatory effect. Thus, state-resolved ChromBPNet models reveal that analyzing autoimmune variants solely in resting B cells would miss this activation-dependent layer of genetic risk. These results underscore the necessity of assembling multi-scale GRNs with chromatin landscapes spanning a continuum of activation and differentiation states to predict and interpret the functional impact of disease-associated non-coding variation on humoral immune responses.

We note that these variant analyses are predictive in nature. Although the predictions are extensively supported by convergent evidence from genetic fine mapping, caQTLs, eQTLs and MPRA assays generated independently of our training data, they do not establish causality at individual loci. Allele-specific enhancer assays and perturbation of individual candidate elements with base editing will be required to convert these prioritized variants into demonstrated causal mechanisms. Therefore, the value of the resource at this stage lies in narrowing large credible sets of variants to a tractable number of state-resolved, experimentally testable candidates.

## RESOURCE AVAILABILITY

### Lead Contact

Further requests and information concerning this study should be addressed to the lead contact, Harinder Singh.

### Materials Availability

No new materials were generated in this study.

### Data and Code Availability

All sequencing data has been deposited to the IGVF Data Portal (https://data.igvf.org/): IGVFDS4024LHFB, IGVFDS7859EYQR, IGVFDS8967LLYA, IGVFDS8822AFTR, IGVFDS9764OTVQ, IGVFDS8419ZMNL, IGVFDS2610IANK, IGVFDS7476HGYD, IGVFDS5867EPVA. Scripts and associated documentation necessary to reproduce the analysis of genomics datasets have been deposited to GitHub: https://github.com/pitt-csi/humanBcell_GRN.

## Acknowledgements

This work was primarily supported by National Institutes of Health (NIH) grant 5U01HG012041 (to N.S., J.D., H.S). This work was also funded by NIH grants 5R01AI170108 (to H.S. and J.D.), 5U01AI179514 (to H.S. and J.D.), UM1HG012076 (to A.T.S.), UM1HG012003 (to. H.W.), 5R01AI130010 (to D.M) and 5T32AI089443 (to N.A.P). N.A.P is a Cancer Research Institute Irvington Fellow supported by the Cancer Research Institute (CRI Award #4185). Flow cytometry and sorting was performed using equipment maintained by the Unified Flow Core (University of Pittsburgh, Department of Immunology). Single-cell capture and sequencing library preparation for Donor 1 experiments was performed by Single Cell Core (University of Pittsburgh, Dr. Robert Lafyatis laboratory). We thank Tracy Tabib and Heidi Monroe for assisting in experimental design, planning and execution of single-cell genomics experiments. Next generation sequencing was performed by either High Throughput Genomics Core (UPMC) (formally the UPMC Genome Center) or Health Sciences Sequencing Core (University of Pittsburgh). This research was supported in part by University of Pittsburgh Center for Research Computing and Data, RRID:SCR_022735, through the resources provided. Specifically, this work used the HTC cluster, which is supported by NIH award number S10OD028483. This work used bridges-2 (EM,RM,GPU) and ocean systems at the Pittsburgh Super Computing Center through allocation CIS240372 from the Advanced Cyberinfrastructure Coordination Ecosystem: Services & Support (ACCESS) program, which is supported by U.S. National Science Foundation grants # 2138259, #2138286, #2138307, #2137603, and #2138296. The Genotype-Tissue Expression (GTEx) Project was supported by the [Common Fund](https://commonfund.nih.gov/GTEx) of the Office of the Director of the National Institutes of Health, and by NCI, NHGRI, NHLBI, NIDA, NIMH, and NINDS. Fine-mapped credible sets for CAP, GENCORD, GEUVADIS, GTEx, and TwinsUK LCL eQTL datasets (QTD000056, QTD000061, QTD000095, QTD000110, QTD000221, QTD000539) were obtained from the eQTL Catalogue (https://www.ebi.ac.uk/eqtl/) on August 7, 2026. We thank Richard James (Seattle Children’s) for guidance on CRISPR/Cas9 editing of primary human B cells. We thank Stephen Yi (Baylor College of Medicine), Mark Shlomchik (University of Pittsburgh), Anshul Kundaje (Stanford), Kyle Ferchen, Nathan Salomonis and Lee Grimes (CCHMC) for general discussions and feedback.

## Author Contributions

Conceptualization, N.A.P., J.F., H.S.; Data Curation, N.A.P, J.F., S.K.; Formal analysis, N.A.P., J.F., S.K., J.S., S.B.G, T.C., A.S., Z.H.R. ; Funding Acquisition, N.A.P., A.T.S., N.S., J.D., H.S.; Investigation, N.A.P, J.F., C.S.M, W.Z., G.M.V, D.B., H.W.; Methodology, N.A.P., J.F., S.K., P.G., A.S., Z.R., D.B., A.K.J., L.L., C.M., D.M., J.D., H.S.; Project Administration, N.A.P., N.S, J.D., H.S.; Resources, H.W., A.T.S, N.S., J.D., H.S.; Software, N.A.P., J.F., S.K., J.S., S.B.G, P.G., A.S., Z.H.R.; Supervision, N.S., J.D., H.S.; Validation, N.A.P., J.F., S.K.; Visualization, N.A.P., J.F., S.K.; Writing – Original Draft, N.A.P., H.S.; Writing – Review & Editing, N.A.P., J.F., S.K., S.B.G, G.M.V, N.S., J.D., H.S.

## Supplementary Information

**Supplementary Table 1**. Single-cell RNA-seq Leiden cluster marker gene lists.

**Supplementary Table 2**. Differential gene expression analysis of BCL6-deficient B cells.

**Supplementary Table 3**. Gene ontology analysis of Donor 1 multiome expression topic genes.

**Supplementary Table 4**. ChIP-seq peak enrichment at Donor 1 multiome accessibility topic peaks.

**Supplementary Table 5**. State-specific GRN TF-gene edges for *in vitro*-differentiated and tonsil B cells.

**Supplementary Table 6**. State-specific GRN TF centrality scores for *in vitro*-differentiated and tonsil B cells.

**Supplementary Table 7.** Perturbation scores from *in silico* simulations for *in vitro* and tonsil B cell GRNs and comparison to prior literature.

**Supplementary Table 8**. Comparison of predicted and experimental effects of TF perturbations on preGC versus PB trajectories.

**Supplementary Table 9**. ChromBPNet model evaluation metrics for held-out chromosomes.

**Supplementary Table 10**. ChromBPNet TF motif family seqlet counts per chromatin accessibility topic.

**Supplementary Table 11**. Normalized TF gene expression across B cell states from Donor 1 multiome data.

**Supplementary Table 12**. RP model scores of seqlet-containing CREs linked to SPIB, BATF, IRF4, and PRDM1 target genes.

**Supplementary Table 13**. Target genes common to IRF4 and its putative binding partners.

**Supplementary Table 14**. Gene ontology analysis of target genes common to IRF4 and its putative binding partners.

**Supplementary Table 15**. Evidence for TF-TF cross-regulation from CRISPR perturbation experiments and comparison to prior literature.

**Supplementary Table 16**. ChromBPNet-predicted variant effects at B cell open chromatin regions across B cell states and non-B cell types.

**Supplementary Table 17**. GM12878 MPRA enhancer analysis of autoimmune-associated variants within B cell open chromatin regions.

**Supplementary Table 18**. Analysis of expression-modulating variants across six different LCL MPRA datasets.

**Supplementary Table 19**. TF motif matches for ChromBPNet-predicted functional variants in each B cell state.

**Supplementary Table 20.** ChromBPNet-predicted functional eQTL variants.

**Supplementary Table 21**. Gene ontology analysis of genes associated with ChromBPNet-predicted functional eQTL variants.

**Supplementary Table 22**. Antibodies used in this study.

**Supplementary Table 23.** Guide RNAs used in this study.

**Supplementary Table 24.** Primers used in this study.

## Methods

### Primary B cell purification and *in vitro* differentiation

Peripheral blood mononuclear cells (PBMCs) from healthy donor leukapheresis products (StemCell Technologies) were isolated by Ficoll density centrifugation. Donors were recruited with institutional review board approval and informed consent by STEMCELL Technologies. Naïve B cells were purified by negative selection using magnetic bead separation (EasySep Human Naïve B Cell Isolation or Memory B Cell Isolation Kits, STEMCELL Technologies). Purified naïve B cells were labeled with either 2µM CFSE for 10 minutes at 37°C or 5µM CellTrace Violet (CTV) for 20 minutes at room temperature before washing with complete RPMI. Naïve B cells were centrifuged and resuspended at 7.5 x 10^5^ cells/mL in the activation media [RPMI-GlutaMAX with 10% heat-inactivated FBS, 1X Penicillin-Streptomycin, 2.6ug/mL of anti-IgM (Jackson ImmunoResearch), 400ng/mL CD40L (Biolegend), 2ng/mL IL-2 (Biolegend), 1ug/mL CpG ODN-2006 (Miltenyi)]. Naïve B cells were cultured in 24-well plates for four days before harvesting and washing with complete RPMI. After washing and centrifugation, activated cells were re-suspended at 4.5 x 10^5^ cells/mL in differentiation media [RPMI-GlutaMAX with 10% heat-inactived FBS, 1X PenStrep, 400ng/mL CD40L (Biolegend), 2ng/mL IL-2 (Biolegend), 5ng/mL IL-4 (Biolegend), 12.5ng/mL IL-10 (Biolegend)]. All centrifugation steps were performed at 300xg for 10 minutes at room temperature.

### Flow cytometry procedures and analysis

Cells were incubated with human Fc block (Biolegend) for 5 minutes at room temperature before staining with cell surface antibodies in FACS buffer [PBS with 2% FBS and 1mM EDTA] with 1X BD Brilliant Stain buffer (BD Biosciences) for 30 minutes on ice. Cells were washed with FACS buffer twice before resuspending in 1X fixation buffer (eBioscience Foxp3 / Transcription Factor Staining Buffer Set) and incubating for 20 minutes at room temperature. Cells were centrifuged at 600xg for 5 minutes before washing with 1X permeabilization buffer (eBioscience Foxp3 / Transcription Factor Staining Buffer Set), centrifuged again and resuspended in 1X permeabilization buffer with 2% rat serum blocking solution. Cells were incubated for 15 minutes at room temperature before adding intracellular antibodies and incubating overnight at 4C. Cells were then washed three times with 1X permeabilization buffer before resuspending in FACS buffer and analyzing on an Attune NxT (Thermo Fisher Scientific) or an Aurora (Cytek). See **Supplementary Table 22** for antibody details.

### Single-cell RNA-sequencing and BCR-sequencing procedure

Naïve B cells from Donor 1 were isolated and activated as described above every two days before harvesting cells from asynchronous cultures on Day 6. Cells were washed with FACS buffer before removing dead cells by magnetic cell isolation (EasySep Dead Cell Removal Kit, STEMCELL Technologies). Live cells were incubated in Human TruStain FcX Fc Receptor Blocking Solution (Biolegend) for 10 minutes on ice before washing and resuspending in PBS with 10% FBS. Cells were then stained with TotalSeq-C anti-human hashtag oligonucleotide conjugated antibodies (Biolegend, see **Supplementary Table 22** for antibody details) for 30 minutes on ice. Cells were then washed three times with 10% FBS PBS before processing with Single-cell 5’ Kit v2 (10x Genomics) and Human BCR Amplification Kit (10x Genomics) according to manufacturer’s instructions. Resulting libraries were sequenced with a NovaSeq 6000 S1-100 cycle flow cell (v1.5) (Illumina).

### Single-cell RNA-sequencing and BCR-sequencing data analysis

FASTQ files were generated using CellRanger (v7.1.0) ‘mkfastq’ function and used as input into CellRanger (v7.1.0) ‘multi’ function using GRCh38 (2020) as reference genome. After demultiplexing using HashSolo^65^, MIRA expression topic modeling was performed using the most highly variable genes. Support Vector Machine (SVM) classification analysis was performed using the scikit-learn Python module^13^. Gene expression values from the Human Tonsil Atlas B cell dataset were standardized using z-score normalization using the ‘StandardScaler’ function. The resulting transformation was applied to the gene expression values from the *in vitro* B cell dataset. C-Support Vector Classification (SVC) was used to fit a model using the scaled Human Tonsil Atlas B cell gene expression matrix as the training dataset. The resulting model was used to generate classification probabilities for the *in vitro* B cell dataset using ‘predict_proba’. Up-regulated tonsil GCBC versus tonsil activated B cell genes were derived by running the Differential Gene Expression module on the Human Tonsil Atlas dataset in the Cell Annotation Platform (https://celltype.info/project/389/dataset/825/md). Gene set enrichment analysis of up-regulated tonsil GCBC genes was performed using the ‘prerank’ function from GSEApy^66^. BCR-sequencing FASTQ files were processed with CellRanger (v7.1.0) ‘multi’ function using GRCh38 (2020) as the reference genome (‘refdata-cellranger-vdj-GRCh38-alts-ensembl-7.0.0’).

### Single-cell multiome procedure

For Donor 1 single-cell multiomics, naïve B cells were isolated and activated as described above every 24 hours before harvesting asynchronous cultures on Day 6. Dead cells were removed by magnetic cell isolation (EasySep Dead Cell Removal (Annexin V) Kit, StemCell Technologies) and stained with fixable viability dye (FVD) (Thermo Fisher Scientific) for 30 minutes on ice before washing with FACS buffer and purifying by fluorescence-activated cell sorting (FACS). See **Supplementary Table 22** for antibody details. FACS-purified cells were washed with 1mL of PBS with 0.04% BSA twice before adding 100µL of chilled 10x Lysis Buffer [10mM Tris-HCl, 10mM NaCl, 3mM MgCl2, 0.1% Tween-20, 0.1% IGEPAL, 0.01% digitonin, 1% BSA, 1mM DTT, 1U/µL RNase inhibitor (Protector RNase inhibitor, Sigma)]. Cells were mixed by pipetting 10 times before incubating on ice for 3 minutes. 1mL of chilled 10x Wash Buffer [10mM Tris-HCl, 10mM NaCl, 3mM MgCl2, 0.1% Tween-20, 1% BSA, 1mM DTT, 1U/µL RNase inhibitor (Protector RNase inhibitor, Sigma)] was added to lysed cells and mixed five times by pipetting. Cells were washed three times with 10x Wash Buffer before resuspending in Diluted Nuclei Buffer [1X Nuclei Buffer (included in 10x Genomics Single-cell Multiome ATAC Kit A), 1mM DTT, 1U/µL RNase inhibitor] for counting and processing with Single-cell Multiome ATAC Kit (10x Genomics). Resulting libraries were sequenced on a NovaSeq SP 100 cycle flow cell (v1.5) (Illumina). For Donor 2 single-cell multiomics, live cells from each timepoint were FACS-purified as described above and were frozen in Bambanker medium (FUJIFILM Irvine Scientific). Upon thawing, nuclei were isolated as described above and were processed according to an adapted version of the MULTI-seq protocol previously described^67^. Briefly, nuclei were resuspended in chilled PBS (750-1000 nuclei/µL) and aliquoted (180 µL per sample). A 1 µM Anchor-Barcode complex (final 50 nM) was added, mixed, and incubated on ice for 5 minutes, followed by the addition of 1 µM Co-Anchor (final 50 nM) with another 5 minutes incubation. Labeling reactions were quenched with 2% BSA in PBS and nuclei were centrifuged at 500xg for 4 minutes at 4°C. After resuspension, samples were pooled and centrifuged again before being resuspended in Diluted Nuclei Buffer for loading and processing according to the Chromium Next GEM Single-cell Multiome ATAC + Gene Expression User Guide (10x Genomics) through the “Step 4: Pre-Amplification PCR step”. 10µL of the ‘Pre-Amplification SPRI Cleanup’ product was fractionated by incubating with 0.6X SPRI beads. The resulting supernatant containing the MULTI-seq barcodes was further purified by SPRI (3.2X) and isopropanol (1.8X). After performing two 80% ethanol washes and elution from SPRI beads, barcode DNA libraries were amplified using Kapa HiFi HotStart Ready Mix with universal I5 primer and TruSeq RPI primers. The resulting PCR product was purified by SPRI-clean up (1.6X), two 80% ethanol washes, and elution in EB Buffer. cDNA (RNA product) and gDNA (ATAC product) libraries were prepared according to Chromium Next GEM Single-cell Multiome ATAC + Gene Expression User Guide.

### Single-cell multiome data analysis

FASTQ files were generated using CellRanger (v7.1.0) ‘mkfastq’ function and used as input into CellRanger ARC pipeline (v2.0.2) using GRCh38 (2020) as the reference genome for alignment, filtering, barcode counting, peak calling and molecule counting using the ‘count’ function. Outputs from each GEM well were aggregated using the ‘aggr’ function. Using SCANPY^68^, cells were filtered with a minimum of 200 genes/cell (with < 20% mitochondrial reads) and genes were filtered with a minimum of 20 cells/gene. Variation due to cell cycle phase was regressed out using the SCANPY^68^ ‘regress_out’ function with G1/S and G2/M gene lists previously described^69^. MIRA expression topic modeling was performed using the most highly variable genes. Following MIRA accessibility topic modeling, a joint embedding space was constructed by joining the transformed expression and accessibility topic spaces. HashSolo^65^ was used for demultiplexing Donor 2 sample timepoints based on MULTI-seq barcode tag levels. Support Vector Machine classification analysis was performed as described above using standardized gene expression matrices from each dataset and applying C-Support Vector Classification and ‘predict_proba’ with the scikit-learn module^13^. Integration of our *in vitro* B cell dataset with that of the Human Tonsil Atlas B cell dataset was performed using the ‘RunHarmony’ function of the harmonypy package^28^.

### STREAM pseudotime and differentiation trajectory inference

Pseudotime inference for B cell differentiation was performed using STREAM. Given the substantial differences in gene expression profiles between naïve B cells and other differentiation states, their inclusion introduced excessive noise in pseudotime inference. Therefore, naïve B cells were excluded from the analysis. Prior to running STREAM, gene expression counts were imputed using Palantir to enhance trajectory reconstruction, with diffusion map dimensionality set to 50 components for the male dataset and 10 components for the female dataset. Highly variable genes were selected by setting the loess fraction to 0.005 and retaining the top 80th percentile of genes for the male dataset, while for the female dataset, the loess fraction was set to 0.01 with the top 90th percentile of genes retained. Dimensionality reduction was performed using modified locally linear embedding (MLLE) for male samples and standard locally linear embedding (SE) for female samples.

Following dimensionality reduction, elastic principal graphs (EPGs) were fitted and optimized to construct branching trajectories. The parameters for fitting the graph were tuned to achieve biologically meaningful differentiation pathways. This approach enabled robust trajectory reconstruction, capturing key differentiation pathways while minimizing artifacts introduced by naïve cells. The resulting STREAM-derived pseudotime values were used to generate a differentiation gradient object using CellOracle’s ‘Gradient_calculator’ module. The resulting gradient object was then used to generate differentiation flow vector field graphs to model normal and perturbed differentiation trajectories.

### MultiVI integration and Moscot optimal transport analysis

A joint latent embedding of the two modalities was learned with MultiVI (scvi-tools v1.4.1), a multimodal variational autoencoder to model gene counts with a zero-inflated negative-binomial likelihood (gene-specific dispersion) and binarized peak accessibility with a Bernoulli likelihood with learned region factors, sharing a low-dimensional Gaussian latent across modalities. The input MuData comprised 28,494 cells passing QC in both modalities, with raw counts for 3,018 highly variable genes and 191,255 ATAC peaks; no batch or covariate keys were supplied. We used the scvi-tools default architecture: two-layer encoders and decoders with 128 hidden units, an 11-dimensional latent space, dropout 0.1, layer normalization, equal modality weights and a Jeffreys modality penalty (75.9M parameters fit to 28,494 × 194,273 ≈ 5.5B observed feature values, ∼73 observations per parameter). To monitor overfitting, training followed the scvi-tools default 90/10 split: the model was fit on 25,645 cells and evaluated each epoch on 2,849 held-out cells, for 200 epochs, with the KL term linearly fitting over the first 50 epochs. The per-cell posterior mean of the 11-dimensional latent was stored as X_MultiVI and defined the feature space for the downstream K-NN graph and optimal-transport (OT) computations. The cell-cell coupling matrix between day 3/4 (source marginal) and day 5/6 (target marginal) cells was computed with <moscot.problems.time.TemporalProblem>. Before solving the OT problem, the marginals were filtered to drop any cell-state contributing fewer than 100 cells to a given temporal-bin, yielding {day3_4: [’ActB-2’, ’ActB-3’, ’ActB-4’, ’PB-1’], day5_6: [‘preGC-1’, ‘preGC-2’, ‘PB-2’]} states per temporal-bin. The default ’Euclidean’ cost used in Waddington-OT, was replaced with a manifold curvature aware, heat-diffusion cost. A kNN graph (k = 30) of the union of day 3/4 and day 5/6 cells in ‘X_MultiVI’ was built as the geodesic geometry via <prob.set_graph_xy(G, t=100.0)>, which implements ’Geodesic Sinkhorn’ by using the heat kernel K_t = exp(−tL) on the graph Laplacian L directly as the Sinkhorn Gibbs kernel. A much larger t = 100 used here (than the default t = ε/4), makes the optimization problem diffused, smoothing local kNN noise so transport mass flows along the global differentiation manifold rather than along short-range Euclidean distances. The problem was solved with entropic Sinkhorn, using parameters <epsilon=1e-3, scale_cost=‘mean’>. From the resulting transport plan T_{day 3/4 -> day 5/6}, we computed the backward (ancestor) cell-state transition matrix shown in Fig. 2h.

### Gene Regulatory Network Inference with MIRA and CellOracle

MIRA regulatory potential (RP) modeling was performed for all highly variable genes with TSS information (4306 genes). The resulting gene-specific RP models were then used in conjunction with CistromeDB-processed ChIP-seq peaks^27^ (6912 datasets) to perform probabilistic *in silico* deletion (pISD) modeling. pISD scores for each TF-gene pair were calculated by removing all peaks that overlap given TF ChIP-seq peak set and measuring the change in the target gene’s RP model. For cases in which multiple ChIP-seq datasets are available for given TF, the top pISD score for each TF-gene pair was retained for downstream analysis.

To perform CellOracle GRN modeling, only TF-gene pairs with a pISD score among the top 5% of all scores were retained as binarized linkages which serve as ‘base-GRN’. This resulted in ∼215 predicted target genes per TF. Only TFs present in the human TF database^37^ were included. For TFs which lack MIRA inferred linkages (due to ChIP-seq availability or quality), we incorporated co-accessibility-based TF-gene linkages inferred using Cicero (co-accessibility > 0.8) from the scATAC-seq data in the multiome. Cicero-derived linkages were added only for target genes present in the highly variable gene set from MIRA for which RP and pISD modeling was performed. The final ‘base GRN’ for CellOracle modeling was constructed as the union of the MIRA-derived TF-gene linkages and Cicero-derived TF-gene linkages so that TFs which lack appropriate ChIP-seq data are not removed or penalized in the downstream predictive modeling steps. CellOracle GRN modeling was performed separately for each of the Leiden clusters in the Donor 1 and Donor 2 multiome datasets. After initially performing fitting using all TF-gene links in base-GRN, the top 10,000 TF-gene linkages (p < 0.001) in each cell state were selected for a second round of fitting using Bayesian ridge regression. The resulting second fitted TF-gene linkages were used for final state-specific GRN assembly, centrality inferences, and *in silico* TF perturbation simulations.

### In silico TF perturbation analysis

For *in silico* TF perturbation analysis, differentiation trajectories as inferred by STREAM pseudotime and CellOracle gradient flow (see ‘STREAM pseudotime and differentiation trajectory inference’) were analyzed along two distinct trajectories: PB (PB-1 and PB-2 clusters) and preGC (preGC-1 and preGC-2 clusters) for in vitro B cell GRNs and PB/PC and GCBC annotations for the tonsil B cell GRNs. For each TF perturbation, the perturbation scores for cells in either PB/PC or preGC/GCBC clusters were computed by calculating the dot product between the reference differentiation flow and the simulated TF perturbation flow. Net perturbation scores for each trajectory were calculated by summing the positive and negative perturbation scores for cells in PB/PC or preGC/GCBC clusters, separately. Then, the overall preGC-PB or GCBC-PB/PC perturbation score was derived by subtracting the net PB or PB/PC perturbation score from the net preGC or GCBC perturbation score for each TF. As with the GRN modeling, this process was conducted separately for male and female datasets.

### Arrayed CRISPR/Cas9 phenotypic screen

Naïve B cells (NBCs) from either Donor 1 or Donor 2 were isolated and activated as described above. After two days of activation, cells were nucleofected with Cas9:sgRNA ribonucleoprotein (RNP) complexes using a 4D-Nucleofector System (Lonza). To generate RNP complexes, 60 picomoles of arrayed sgRNAs (three per gene/reaction, see **Supplementary Table 23** for sequences) (Synthego/EditCo) were incubated with 20 picomoles of Cas9 (IDT) (diluted in P3 nucleofection Solution with supplement (Lonza)) each for up to an hour at room temperature. Prior to nucleofection, cells were washed with PBS and centrifuged at 90xg for 10 minutes before resuspending in P3 Nucleofection Solution with supplement. 200,000-300,000 cells per reaction were nucleofected in Nucleocuvette strips and re-cultured in activation media as described above. At Day 4, 50,000 cells from each sample were split off for DNA extraction with 50µL of Quick Extract Solution (Biosearch Technologies). Cells were lysed at 68C for 15 minutes followed by 95C for 10 minutes. Primers flanking the most distal pair of sgRNAs for each gene were used to detect deletions by PCR. At Day 6, cells were harvested for flow cytometry analysis as described above. See **Supplementary Table 22** for antibodies used.

### ChromBPNet data analysis

Aggregated cell state-specific ATAC-sequencing reads were extracted from the single-cell multiome dataset by using Sinto’s ‘filterbarcodes’ function. Resulting BAM files were used for peak calling with MACS2 ‘callpeak’ function with the following options: --shift -75 --ext 150 -- pvalue 0.01 --nomodel -B --SPMR. Peaks for each cell cluster were merged to generate a union set of ATAC peaks which were used to train ChromBPNet models for each cell cluster. A Tn5 bias model was trained by running the ChromBPNet ‘bias pipeline’ with nonpeak sequences from inaccessible chromatin regions. Bias-corrected BigWig files were generated by running ChromBPNet ‘pred_bw’ and base-pair resolution contribution score predictions were generated by running ChromBPNet ‘contribs_bw’ with the union set of ATAC peaks. The resulting bigWig files were converted to bedGraph files using bigWigToBedGraph (ucsc_utilities). Profile contribution score tracks (i.e. predicted contribution to peak profile/shape) were used for all downstream analysis of ChromBPNet models. For accessibility topic TFModisco analysis, ‘contribs_bw’ was also run with the top 10,000 peaks for each accessibility topic. The resulting profile-score AnnData objects were used as input into the TFModisco ‘motifs’ function using sampling of 10,000 seqlets and the ‘report’ function using the Jaspar 2024 Core Vertebrates (non-redundant) PFM collection. Seqlets matching motifs in the same Jaspar motif cluster family were merged for quantification of TF motif family seqlet instances across accessibility topics. Homer or CEseek were used to scan for specific single motif instances or composite element motif instances across the union set of peaks or accessibility topic peaks. Aggregated histogram plots were generated using the ‘plotProfile’ function from deepTools.

### CRISPR/Cas9 perturbation with single-cell RNA-sequencing procedure

Naïve B cells from Donor 1 were isolated and activated as described above. After two days of activation, cells were nucleofected with Cas9:sgRNA ribonucleoprotein (RNP) complexes as described above. After nucleofection, cells were re-cultured in activation media for two days. On Day 4, 50,000 cells per sample were separated for PCR genotyping as described above. 200,000 cells per sample were re-cultured in differentiation media for two days and frozen with Bambanker medium and stored at -80C. For BATF, IRF4, PRDM1 and SPIB perturbations, 200,000 cells per sample were also frozen with Bambanker medium and stored at -80C on Day 4. Cells from all timepoints were thawed and dead cells were removed by magnetic cell isolation (EasySep Dead Cell Removal (Annexin V) Kit, StemCell Technologies). Live cells were incubated in Human TruStain FcX (Biolegend) for 10 minutes on ice before washing and resuspending in FACS buffer. Cells were then stained with TotalSeq-C anti-human hashtag oligonucleotide conjugated antibodies (Biolegend, see **Supplementary Table 22** for antibody details) for 30 minutes on ice. Cells were then washed two times with 2% FBS/PBS and one time with 1% BSA/PBS before processing with Single-cell 5’ Kit v2 (10x Genomics) and Human BCR Amplification Kit (10x Genomics) according to manufacturer instructions.

### CRISPR/Cas9 perturbation with single-cell RNA-sequencing data analysis

FASTQ files were generated using CellRanger (v7.1.0) ‘mkfastq’ function and used as input into CellRanger ‘multi’ function using GRCh38 (2024) as the reference genome. CellRanger-demultiplexed sample feature barcode matrices were merged and MIRA (v2.1.0) expression topic modeling was performed using the most highly variable genes. Following Leiden clustering, differential gene expression analysis between knockout and control cells within each cluster was performed using ‘rank_genes_groups’ (method = ‘t-test_overestim_var’) from SCANPY. Genes with adjusted p-values < 0.05 and absolute fold change > 1.2 in each of the two replicates were considered differentially expressed. Energy distance calculations were performed using the ‘Distance’ function from pertpy^20^. Comprehensive gene set enrichment analysis of differentially expressed genes was performed using Metascape [http://metascape.org]^70^. Curated gene set enrichment analysis of MYC target genes^69^ was performed using the ‘prerank’ function from GSEApy^66^.

### Autoimmune disease variant analysis

Fine-mapped GWAS SNPs were downloaded from PICS2^47^ and filtered to retain linked SNPs that were associated with at least one of 28 different autoimmune associated diseases. The variant-scorer script (https://github.com/kundajelab/variant-scorer) was used to compute a predicted effect size of each SNP on local chromatin accessibility using each B cell state-specific ChromBPNet model. Variants with a log2 fold change p-value <0.05 were considered significant. To identify TF motifs at variants, ChromBPNet-derived CWMs were converted into position probability matrices (PPMs) via the Softmax function, ensuring that the probabilities at each position summed to 1. To assess similarity to known TF motifs, the resulting PPMs were compared to each Jaspar 2024 core vertebrate motif PWM via Tomtom. A threshold of FDR < 0.001 was employed to define significant similarity. To link variants to genes, variants were mapped to peaks contained in the feature linkage matrix for the Donor 1 multiome dataset derived from the 10x ARC pipeline which only retains significant linkages (FDR-adjusted p-value < 10^-5^).

### Massively parallel reporter assay (MPRA) and analysis

150 base-pair oligos were synthesized and PCR amplified before adding random barcodes via PCR as previously described^71^. The resulting amplified library with barcodes was cloned into the pJD-Donor plasmid via SpeI/MluI digestion and ligation. The library and barcode regions were amplified and sequenced to assign barcodes to each oligo as previously described^71^. The minimal promoter and luciferase sequence was digested from the pMPRAdonor2 plasmid (Addgene: #49353) using KpnI and XbaI and inserted between the oligo and barcode via KpnI and XbaI digestion and ligation.

GM12878 cells (Coriell Institute for Medical Research) were cultured in RPMI medium (Corning, 10-040-CV) supplemented with 15% FBS (Gibco A52568-01), 1% GlutaMAX (Gibco 35050-061) and 1% Pen-Strep (Corning, 30-002-CI). For each transfection, 1×10⁷ cells were mixed with 10 μg of the TF-motif MPRA library in 100 μl of RPMI. These cells were transfected with a Neon transfection system (Thermo Fisher Scientific, MPK5000) using a kit (MPK10096B) with 3 pulses of 1200 V for 20 ms. A total of five biological replicates were performed, each using cells at a density of ∼1 million cells/ml for the transfections. For each replicate, a total of 1.5×10⁸ cells were pelleted at 200 × g and resuspended in 1.5 ml of RPMI medium containing 150 μg of the TF motif library. After transfection, each replicate was recovered at a density of 5×10⁵ cells/ml in 300 ml of RPMI medium containing 15% FBS, 1% GlutaMAX and 1% Pen-Strep. After 24 hours, the cells were pelleted at 200 × g and washed once with PBS. The cells were resuspended in 15 ml of RLT buffer provided with Qiagen Maxi RNeasy (Qiagen, 75162) and 300 mM DTT (G-Biosciences, 786227). The cells were then homogenized with a tissue homogenizer, TH (OMNI International), for 1 min at maximum speed and the lysates were stored at –80°C.

The resulting RNA was reverse transcribed using the primer Lib_Hand_RT and SuperScript IV reverse transcriptase kit (see **Supplementary Table 24** for primer sequences). The resulting cDNA library was amplified using Luc-F and Luc-R before adding P5 and P7 adapters and indices via a second PCR amplification with P5_F and P7_index_R primers. After SPRI clean-up with AMPure XP beads, the resulting library was sequenced with the custom R1_sequencing_primer and index_sequencing_primer using a NextSeq 2000.

To process the MPRA data, we employed an established pipeline (https://github.com/tewhey-lab/MPRA_oligo_barcode_pipeline). The barcode-oligo sequencing FASTQ files were parsed using the MPRAmatch function to create the barcode-oligo dictionary, retaining only barcode-oligo links with exact matches. Barcode count tables were subsequently generated using the MPRAcount function. To determine which oligos exhibit enhancer activity and which variants modulate enhancer activity (emVars), we employed an established pipeline called mpraprofiler (https://github.com/WeirauchLab/mpraprofiler)^72^ (see **Supplementary Table 17**). Oligos exhibiting significant (padj < 0.05) positive cDNA:RNA ratios were further analyzed for allelic skewing by comparing the alternative:reference allele counts. Variants with an allelic skew FDR < 0.1 were categorized as emVars for downstream analysis. See Supplementary Table 18 for previous studies that were included for ChromBPNet-predicted functional variant enrichment analysis.

### Quantitative trait loci analysis

Primary B cell caQTL summary statistics from PBMCs^48^ were used to evaluate the concordance between ChromBPNet-predicted and experimentally observed chromatin accessibility effects. caQTL variant coordinates were lifted from hg19 to hg38 using the UCSC hg19ToHg38.over.chain (rtracklayer) file before intersecting with ChromBPNet variant predictions. For variants tested against multiple peaks, the caQTL record with the lowest q-value was retained. caQTL allelic effect sizes were transformed to a log-odds scale (log2(effect/(1-effect))) for comparison with ChromBPNet-predicted log fold change values. Concordance was assessed for variants that were significant caQTLs (q < 0.05) and significant ChromBPNet-predicted variants (p < 0.05). Directional concordance was defined as agreement in sign between predicted log fold change and the caQTL log-odds effect.

Lymphoblast cell line (LCL) caQTL results from 10 different ancestries were also used to evaluate the concordance between ChromBPNet-predicted and experimentally observed chromatin accessibility effects. caQTL results from each ancestry cohort were lifted from hg19 to hg38 using the UCSC hg19ToHg38.over.chain file (rtracklayer). Per-variant significance was combined across cohorts using Fisher’s method and variants were retained if they had p < 0.005 in at least two ancestry cohorts. For each B cell state and range of Fisher’s combined p-values, the ChromBPNet-predicted higher-accessibility allele (p < 0.05) was compared to the caQTL- reported higher-accessibility allele. Directional concordance was tested against a null of 50% using one-sided exact binomial tests.

Fine-mapped cis-eQTL variants from LCLs were used to assess whether ChromBPNet-predicted functional variants are supported by association with changes in gene expression. SuSiE credible-set outputs from six LCL datasets (QTD000056, QTD000061, QTD000095, QTD000110, QTD000221, QTD000539) were obtained from the eQTL Catalogue and combined. Only variants that were assigned to at least one credible set and associated with genes with a median TPM > 1 in the corresponding dataset were retained for downstream analysis. For each variant-gene pair, the dataset with the maximum posterior inclusion probability (PIP) was retained for downstream analysis. eQTL target genes were mapped to gene symbols using org.Hs.eg.db and biomaRt. Gene ontology pathway analysis of ChromBPNet-predicted functional eQTL target genes was performed using Metascape [http://metascape.org]^70^.

---

**Extended Data Fig. 1.**
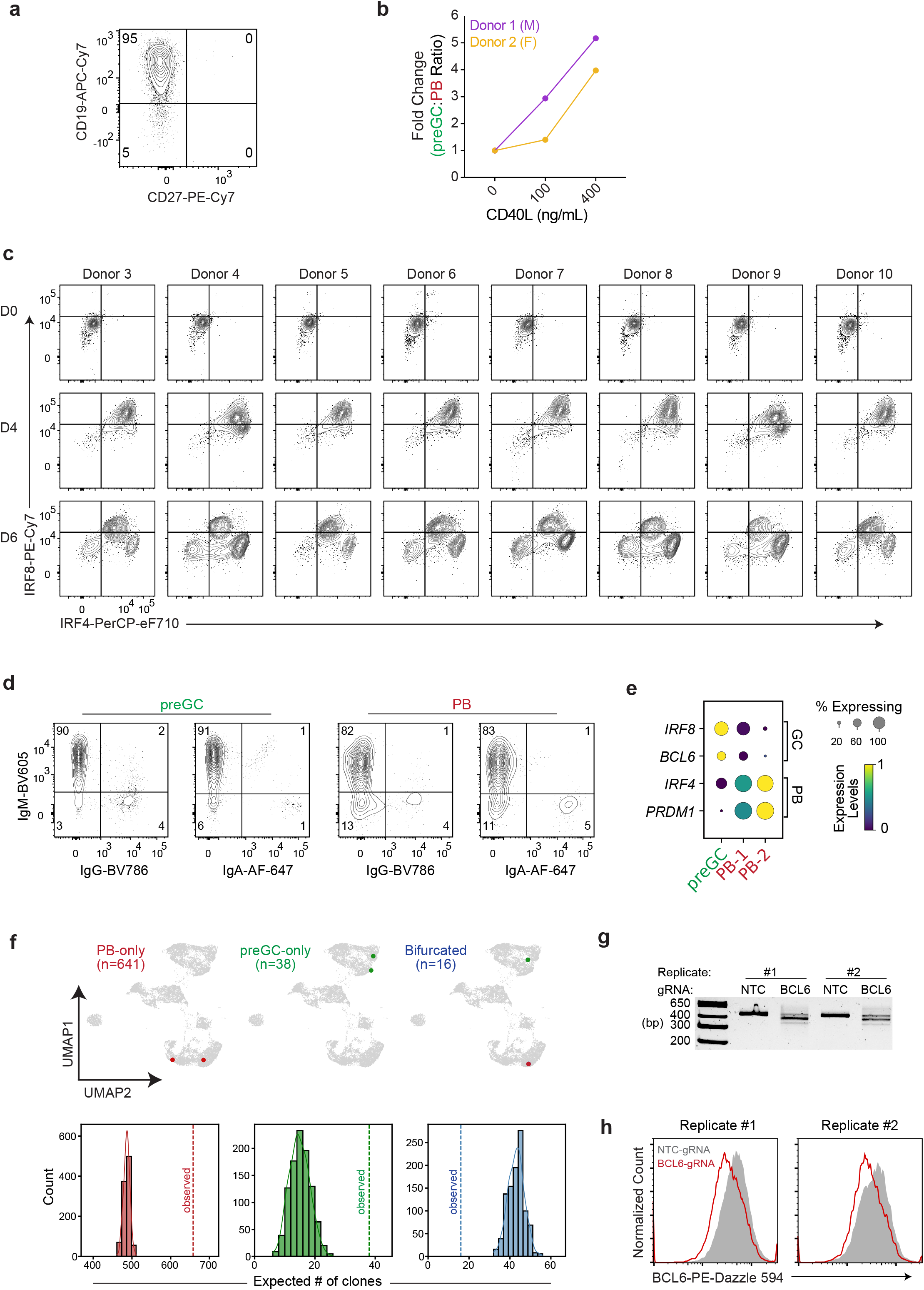
Human B cells activated *in vitro* bifurcate into PB and preGC trajectories. **(a)** Representative flow cytometry analysis of CD19 and CD27 in D0 cells after naïve B cell enrichment. **(b)** Effect of CD40L concentration on generation of preGC vs. PBs. Naïve B cells were stimulated for 4 days as in Fig. 1a, before re-culturing with the indicated concentrations of CD40L in the presence of IL-2, IL-4 and IL-10 for 2 days. preGC (IRF8^hi^IRF4^lo^) to PB (IRF8^lo^IRF4^hi^) ratios were quantified at D6 for two donors (male and female). **(c)** IRF4 and IRF8 expression dynamics in activated B cells from 8 additional donors (4 male and 4 female) by flow cytometry, as in Fig. 1a. **(d)** Representative flow cytometry analysis of IgM, IgG and IgA in preGC and PB compartments at D6. **(e)** Expression of key transcription factor genes across preGC and PB clusters at D6. Dot size represents the percentage of cells expressing each transcript and color intensity indicates mean expression level (standardized between 0 and 1). **(f)** Single-cell BCR-seq analysis of B cell clones at D6. UMAP displaying representative B cell clones colored by fate (top panel). Number of clonotypes (size >1) in each clonal fate category is displayed. Expected versus observed number of unique clones (of size >= 2) per indicated clonotype (bottom panel). Distribution of expected clone numbers was derived using Monte Carlo simulations based on a Poisson distribution of observed clone sizes. (**g**) PCR genotyping results confirming CRISPR editing activity at the BCL6 locus in control and BCL6-KO cells from both donors. Primers flanking the two most distal gRNA target sequences were used to detect large indels. (**h**) Flow cytometry results confirming decreased BCL6 protein levels at Day 6 of culture.

**Extended Data Fig. 2.**
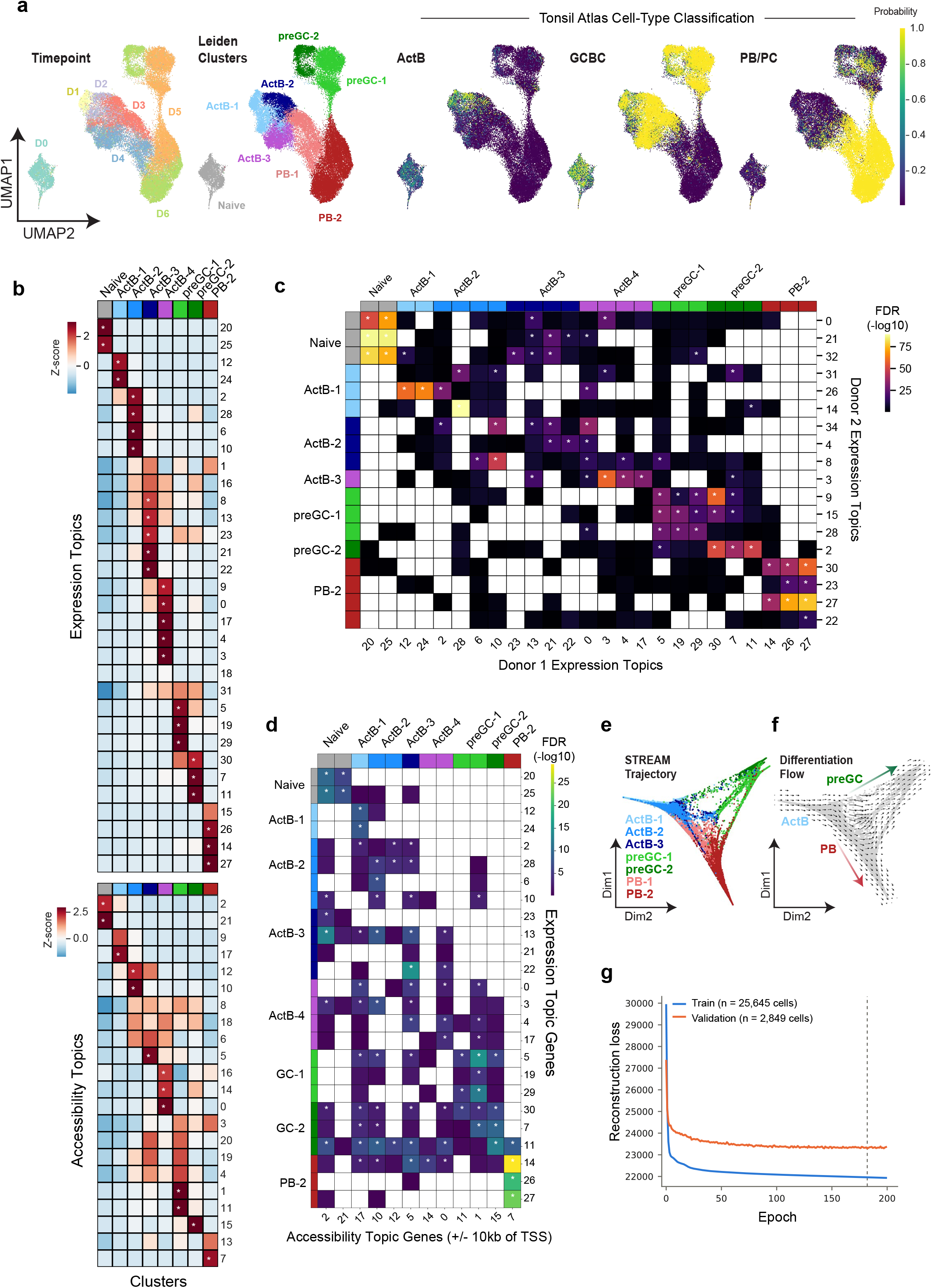
Temporal profiling of chromatin and gene expression dynamics in bifurcating B cell trajectories. **(a)** Naïve human B cells from Donor 2 (female) were activated and differentiated *in vitro* as in Fig. 1a and profiled every 24 hours by paired single-cell RNA/ATAC-seq. MIRA joint topic modeling of transcripts and chromatin accessibility features was used to project cells in low-dimensional UMAP space and colored based on timepoint (*left*), Leiden cluster annotations (*middle*), and support vector machine classification probabilities based on Human Tonsil Atlas cell-types (*right*). **(b)** Heatmaps displaying z-scores of gene expression topics (*top*) or chromatin topics (*bottom*) in the indicated cell clusters. Mean composition values of each topic (rows) in each cell cluster (columns) were used to generate the heatmap. Asterisks indicate topics associated with a particular cell state (z-scores > 2). **(c)** Heatmap displaying the significance of overlapping expression topic gene sets (top 250 genes/topic) in the annotated cell clusters derived from multiome datasets of Donor 1 and Donor 2. False Discovery Rates (FDRs) were derived from Fisher’s exact tests with Bonferroni multiple-comparisons test correction. Asterisks denote highly significant (FDR < 0.001) intersection of gene sets in the indicated expression topic for the designated cell clusters. **(d)** Heatmap displaying the significance of intersection between the top 250 genes in each Donor 1 expression topic and genes linked to top 10,000 peaks in each Donor 1 accessibility topic (+/- 10 kb of gene TSS). FDRs were derived from Fisher’s exact tests with Bonferroni multiple-comparisons test correction. Asterisks denote highly significant (FDR < 0.001) intersection of genes linked to a chromatin topic with genes in the indicated expression topic for the designated cell clusters. **(e)** STREAM trajectory analysis of *in vitro* differentiated B cells, colored by their annotations from Extended Data Fig. 2a (Donor 2). **(f)** Differentiation flow vectors are displayed on STREAM trajectories as in Fig. 2g. **(g)** Line plot displaying training and validation reconstruction loss over 200 epochs for the MultiVI joint model, showing a stable loss gap between training and validation cells with minimal overfitting.

**Extended Data Fig. 3.**
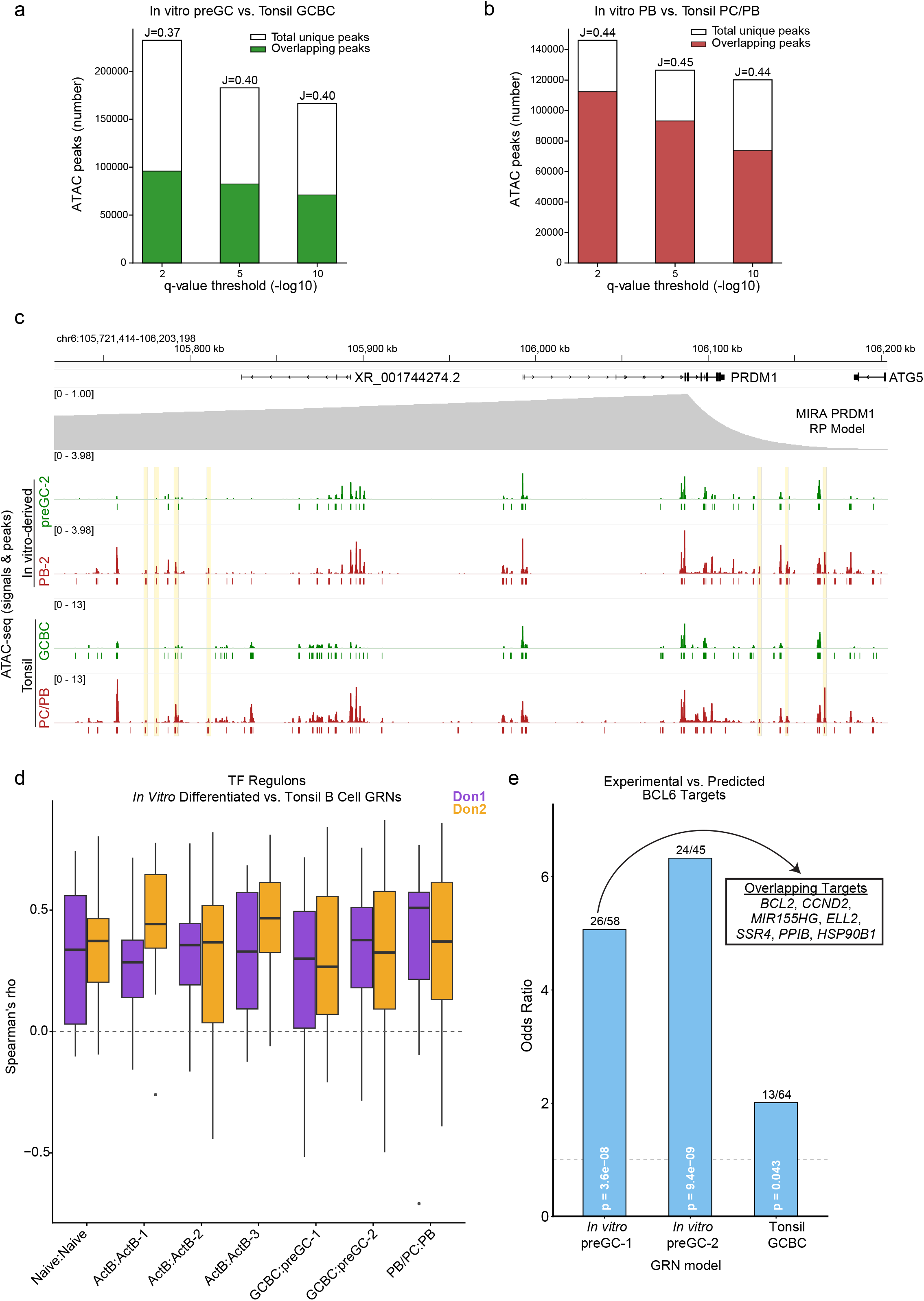
Comparison of *in vitro* differentiated and tonsil B cell GRNs. (**a**,**b**) Intersection of ATAC peaks from *in vitro*-derived B cells and tonsil B cells. (**a**) Bar chart displays the total number of unique peaks passing each q threshold (filled white) and number of overlapping peaks (>= 50bp) (filled green) between *in vitro*-derived preGC states (preGC-1 and preGC-2 clusters combined) and the tonsil GCBC state. Each bar is annotated with the Jaccard similarity index (*J*, scaled 0-1). (**b**) Bar chart displays the total number of unique peaks passing each q threshold (filled white) and number of overlapping peaks (>= 50bp) (filled red) between *in vitro*-derived PB state (PB-2 cluster) and the tonsil PB/PC state. Each bar is annotated with the Jaccard similarity index (*J*, scaled 0-1). (**c**) Representative genome browser view displaying the overlap between *in vitro*-derived and tonsil B cell pseudobulk ATAC-seq peaks. Lines below ATAC-seq signal tracks indicate peaks called at -log10 q > 5. Yellow boxes highlight peaks that were specific to *in vitro*-derived PB and tonsil PB/PC states. (**d**) Box plots display the distribution of Spearman rank correlation coefficients measured by comparing the TF-gene edge weights between *in vitro*-derived and tonsil B cell TF regulons. Comparisons were restricted to TFs with at least 5 shared predicted target genes for each comparison. In vitro-derived GRNs from Donor 1 (male) and Donor 2 (female) are indicated by purple and orange bars, respectively. (**e**) BCL6- KO DEG enrichment among predicted BCL6 target genes. Bar plot displays odds ratio from Fisher’s exact test comparing the overlap between BCL6-KO up-DEGs from Fig. 1h (Day 6 preGC state) with predicted repressed target genes within the indicated GRN model. Numbers above bars indicate number of genes that were BCL6-KO up-DEGs and predicted repressed targets over the total number of predicted repressed targets.

**Extended Data Fig. 4.**
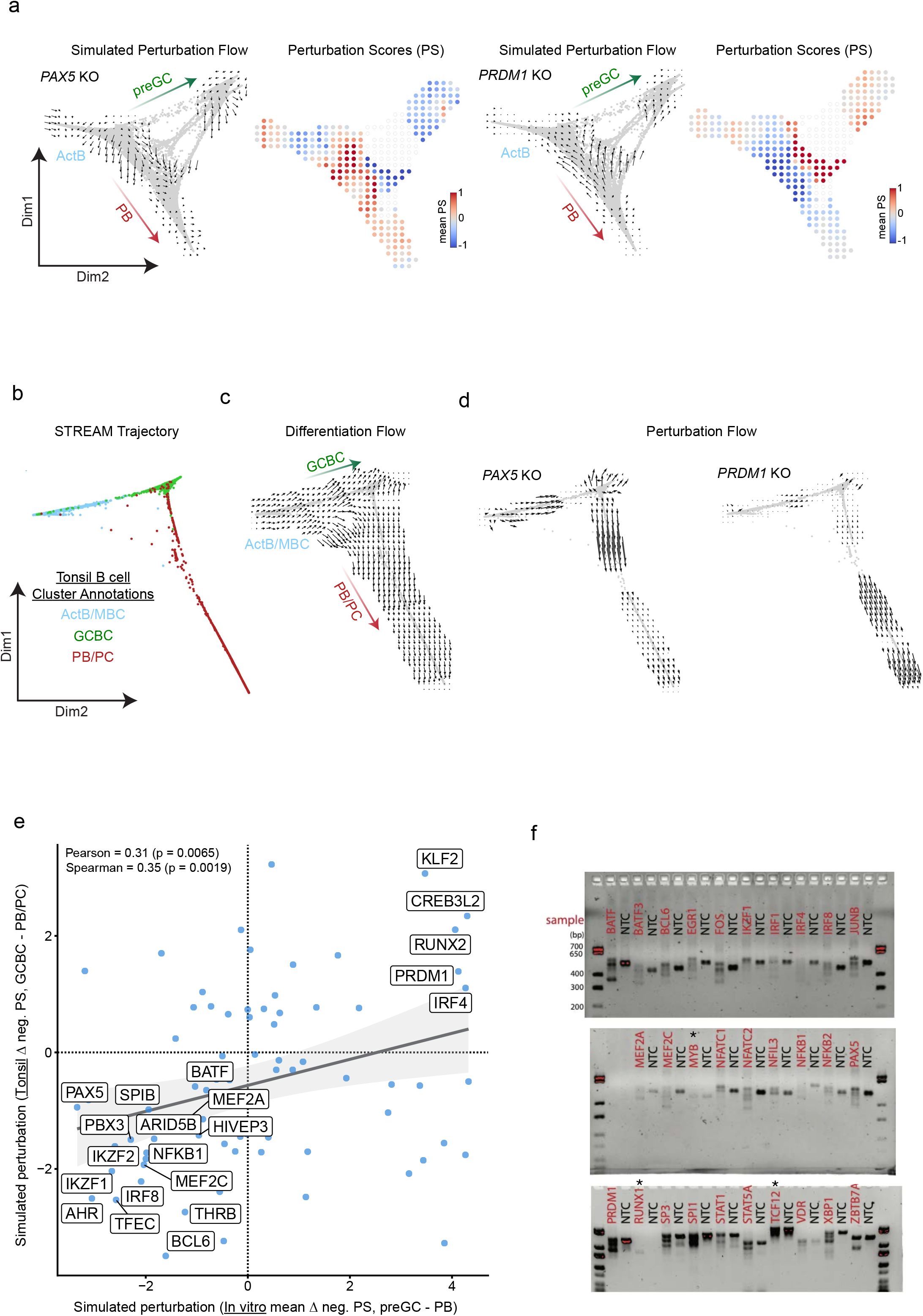
Predicting and testing TF perturbation effects on preGC versus PB trajectories and comparison to tonsil B cell predictions. (**a**) Representative *in silico* TF KO simulations visualized with perturbation flow vector fields or perturbation score grids for PAX5 (*left*) or PRDM1 (*right*) (Donor 2). Perturbation scores (PS) are the dot product of differentiation flow vectors and perturbation simulation vectors. Net preGC and net PB perturbation scores (PS) are derived from summing the PS values for cells in preGC or PB states, respectively. **(b)** STREAM trajectory analysis of tonsil B cells from Human Tonsil Atlas multiome data (see Fig. 2c), colored by their joint profile cell cluster annotations (see Methods). **(c)** STREAM pseudotime values were used with the CellOracle gradient module to generate differentiation flow vector fields (see Methods). (**d**) Representative *in silico* TF KO simulation in tonsil B cells visualized with perturbation flow vector fields for PAX5 and PRDM1 KO. (**e**) Comparison of simulated perturbation results between tonsil and *in vitro*-derived B cell GRN models. The delta negative perturbation score is defined by the difference between negative perturbation scores for cells in GCBC vs. PB/PC (tonsil) or preGC vs. PB (*in vitro*) clusters. Select TFs with concordant simulated perturbation results between tonsil and *in vitro*-derived B cell GRNs are annotated. Gray line and shaded band indicate linear regression fit and 95% confidence interval, respectively. (**f**) PCR genotyping results confirming CRISPR editing activity at indicated TF genes. Primers flanking the two most distal gRNA target sequences were used to detect indels, which appear as shifted or absent bands. Asterisks above indicate samples that had ambiguous or unconfirmed indel formation.

**Extended Data Fig. 5.**
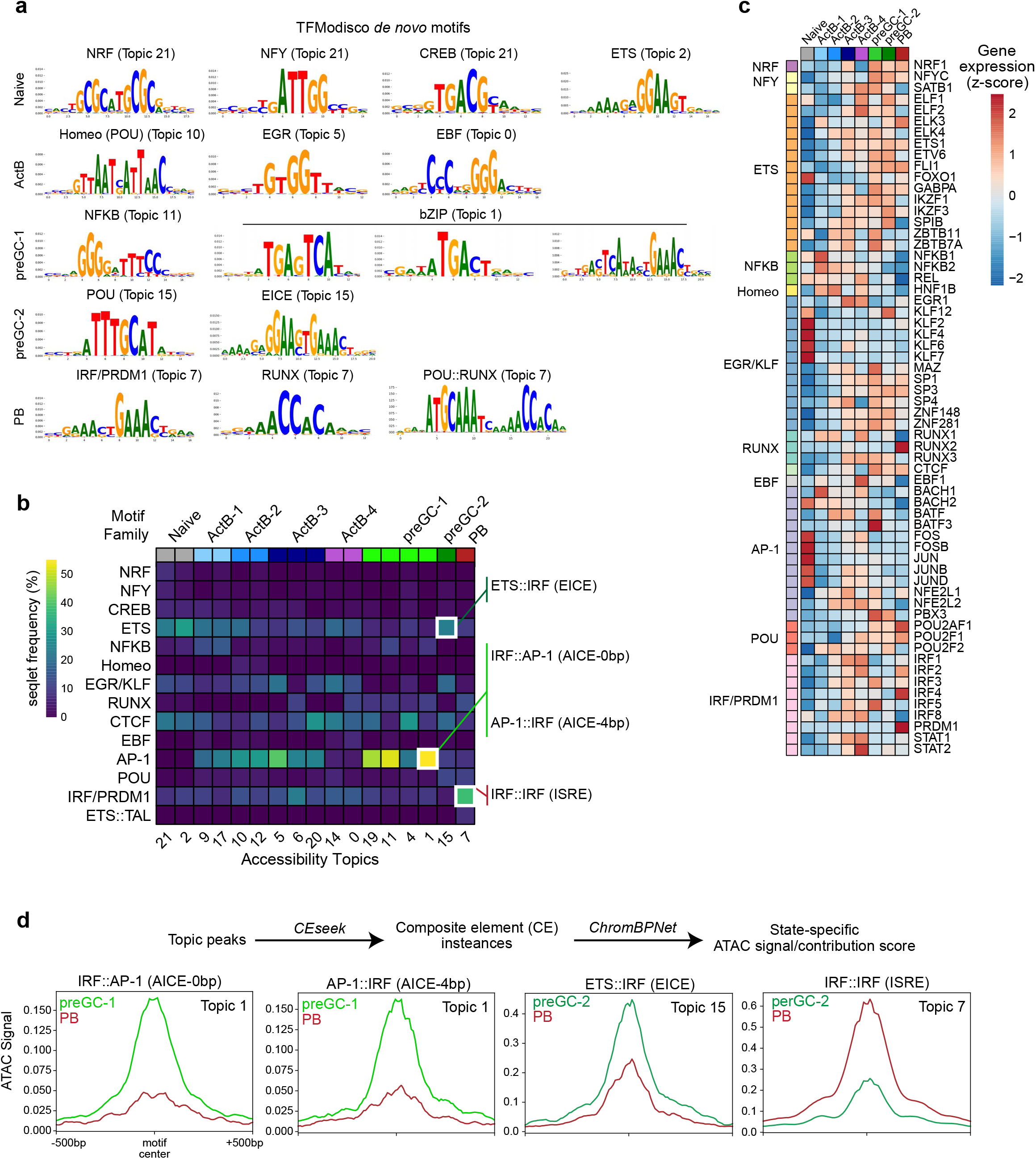
ChromBPNet uncovers dominant and reciprocal TF action at IRF motifs during B cell fate specification. **(a)** Representative *de novo* TF motifs identified by TFModisco are displayed for indicated accessibility topics with their associated cell states indicated on the left. **(b)** Heatmap displaying the raw TFModisco-derived seqlet frequencies for each TF motif family displayed in Fig. 5d. **(c)** Heatmap displaying RNA expression of genes encoding indicated TF family members associated with a given TF motif across various B cell states. **(d)** Aggregated ATAC-seq signals are displayed for preGC or PB cell states at indicated topic peaks containing IRF-related composite elements (EICE, AICE) or IRF::IRF motifs (ISRE).

**Extended Data Fig. 6.**
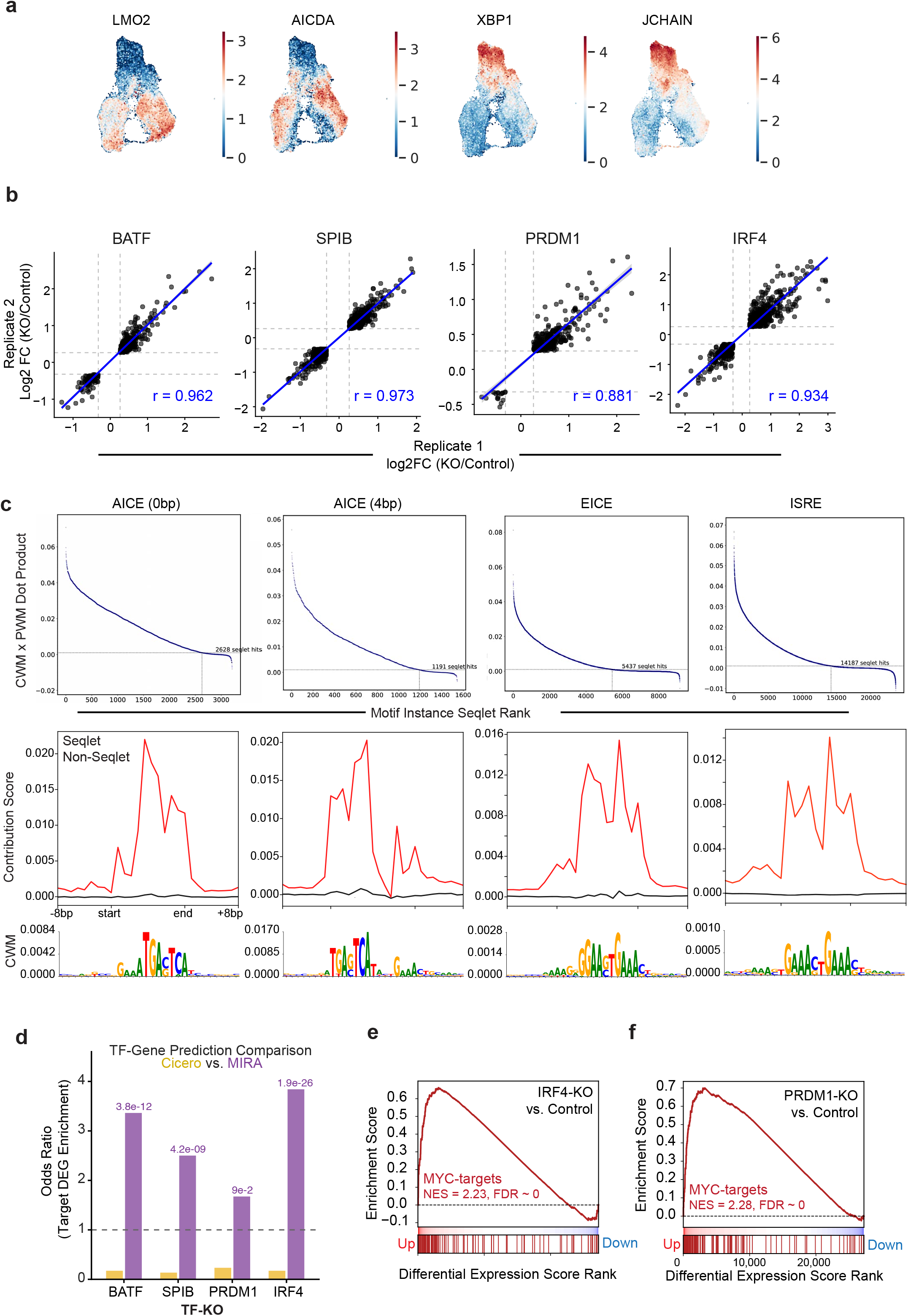
Single-cell perturbation analysis of counteracting PB and preGC TFs. **(a)** UMAPs displaying preGC and PB marker genes using the aggregated scRNA-seq dataset (see Fig. 6a). **(b)** Plots displaying reproducibility between experimental replicates (log2 fold change (FC) in gene expression) for each TF condition (r = Pearson correlation coefficient). Only genes that are statistically significant (adj. p < 0.05) in both replicates are shown, irrespective of directionality. Gray lines indicate |FC| of 1.2. **(c)** Genome-wide ChromBPNet contribution score seqlet-analyses for IRF-containing composite elements or simple IRF motifs in the ActB-3 cell state. Motif instances are displayed in descending order based on their CWM x PWM dot products (*top*). Dot products were normalized to their respective motif lengths. Dotted lines indicate the threshold (0.01) set for calling a motif instance as a seqlet hit. Aggregated contribution score signals at seqlet versus non-seqlet motif instances with CWM logo for seqlet motif instances are shown below each set of panels (*bottom*). (**d**) Comparison of TF-gene linkage prediction methods based on data in Fig. 6c. Odds ratios from Fisher’s exact tests measuring enrichment of TF-KO DEGs within Cicero-based (gray) versus MIRA-based (red) TF-gene prediction sets. P-values for MIRA TF-gene enrichment are shown above bars. (**e**,**f**) Gene set enrichment analysis of MYC target genes in IRF4-KO (**e**) or PRDM1-KO (**f**) cells at D4.

**Extended Data Fig. 7.**
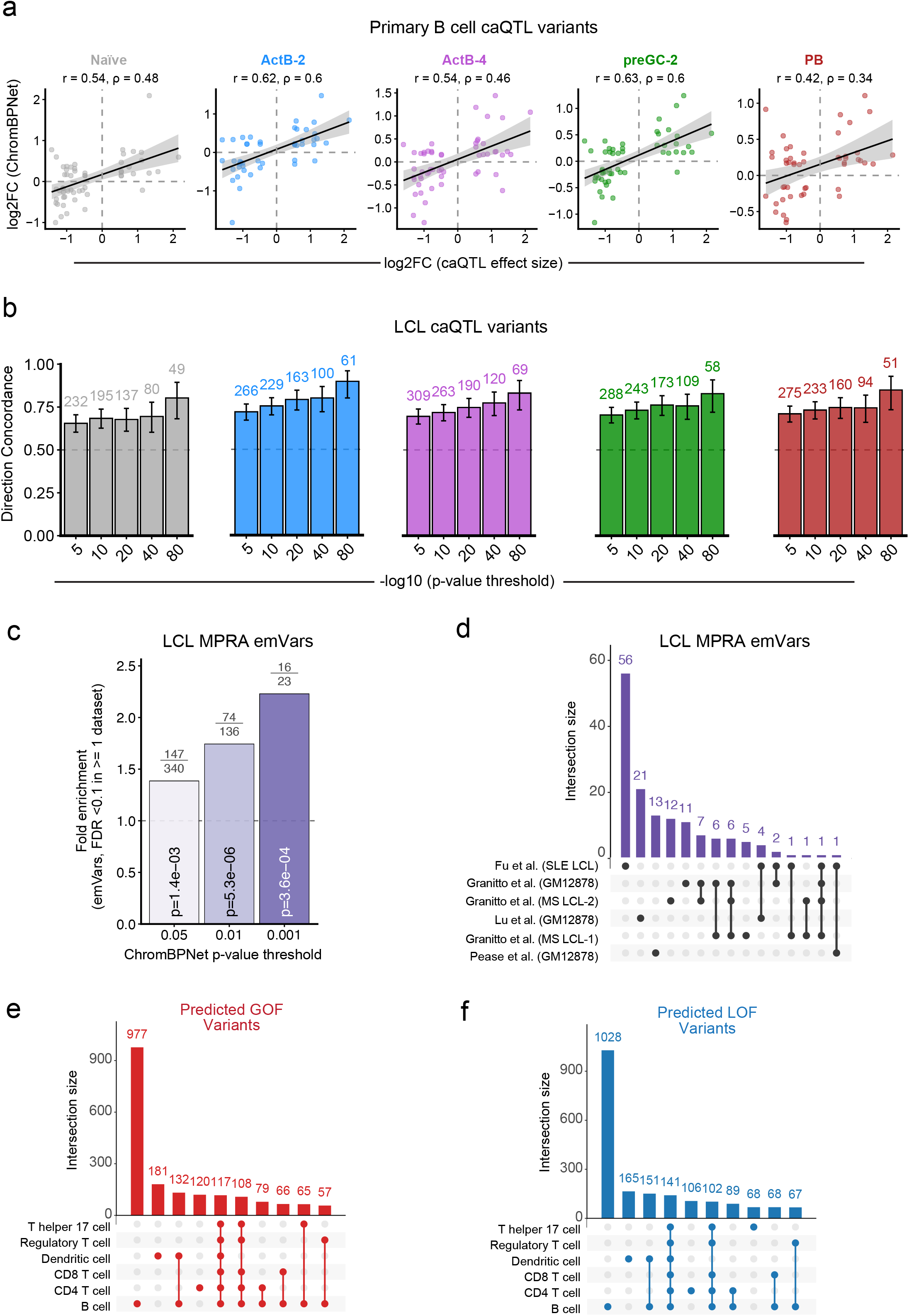
Concordance between ChromBPNet-predicted variant effects and caQTL and MPRA allelic effects, and B cell specificity of predicted variant effects. (**a**) Correlation between predicted and observed chromatin accessibility effects among significant caQTLs reported in total B cells from peripheral blood. Variants that satisfied both thresholds (q < 0.05 (caQTL) and p < 0.05 (ChromBPNet)) are displayed. Pearson’s and Spearman’s rank correlations are displayed above each plot along with a linear regression line with shaded region indicating the 95% confidence intervals. (**b**) Directional concordance between predicted and observed chromatin accessibility effects (i.e. either increased/decreased with alternative allele) among LCL caQTLs. The directional concordance was measured for all ChromBPNet functional variants (p < 0.05) at varying caQTL p-value thresholds. The numbers above bar plots indicate the number of ChromBPNet-predicted functional variants that have directional concordance with the reported caQTL effect at each caQTL p-value threshold. P-values were derived from performing a combined Fisher’s test across all LCL ancestries reported. (**c**) Enrichment of ChromBPNet-predicted functional variants among LCL MPRA-derived expression-modulated variants (emVars). Fisher’s exact test was performed at varying ChromBPNet p-value thresholds, using all tested variants that were non-emVars (FDR > 0.1) and all ChromBPNet variants with p > 0.05 as the background sets. Numbers above plot indicate the number of variants that satisfy both the emVar and ChromBPNet significance thresholds over the total number of variants at each ChromBPNet threshold. The minimum ChromBPNet p-value across all 8 B cell states was used in all calculations. (**d**) Upset plot displaying overlap between predicted functional emVars (ChromBPNet p < 0.05 and emVar FDR < 0.1) derived from different datasets. (**e**, **f**) Upset plots displaying the number of shared and cell-type specific predicted GOF (**e**) or LOF (**f**) variants (p < 0.05). The top 10 most numerous intersection sets are displayed. B cell variants are represented by the prediction from the B cell state with the smallest p-value. Non-B cell datasets were derived from *in vitro* activated samples (ENCODE Portal: ENCSR115OIV, ENCSR261PWP, ENCSR222SCA, ENCSR863TVR, ENCSR373NFA).

**Extended Data Fig. 8.**
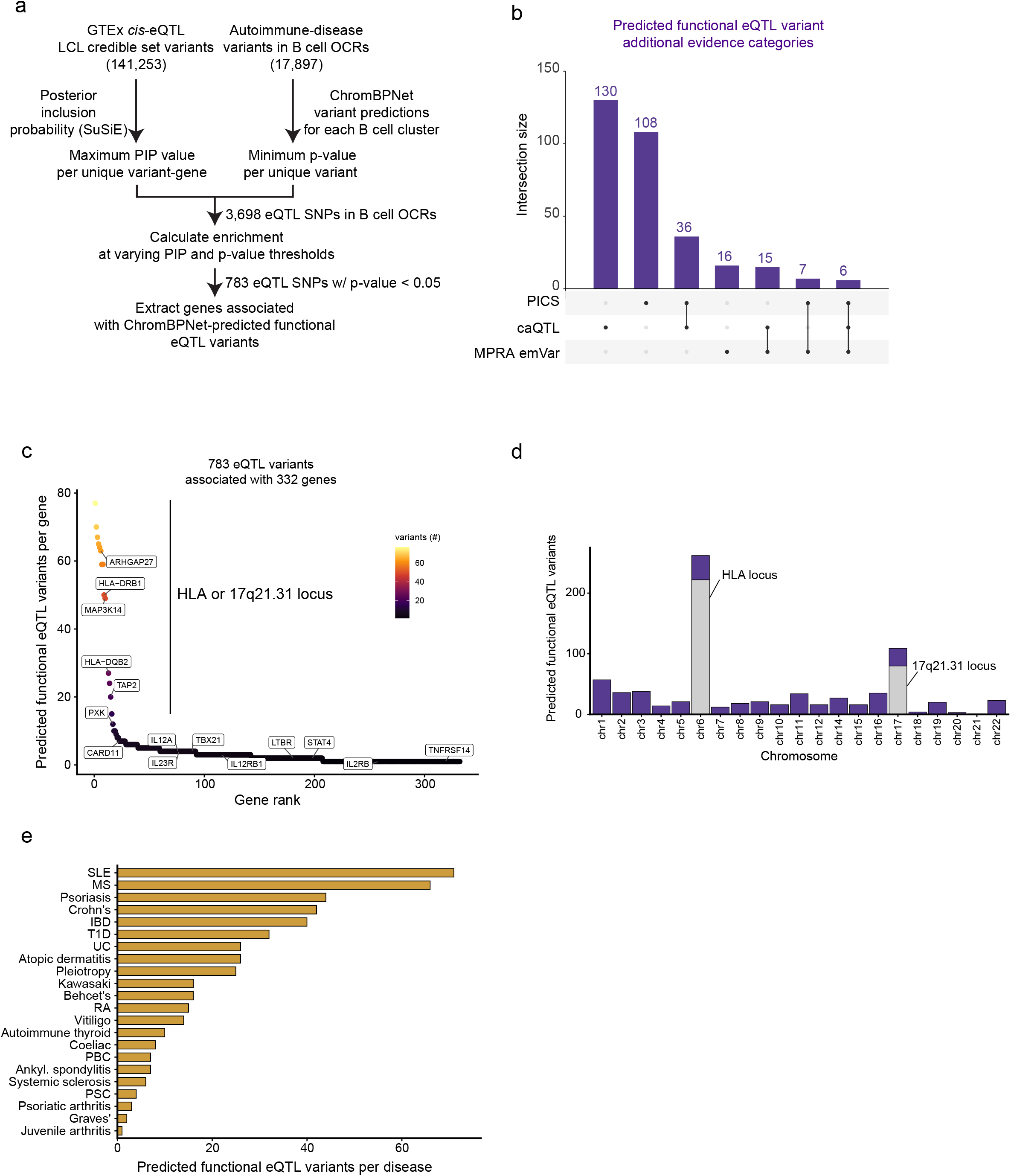
Predicted functional variants are enriched for eQTL variants linked to autoimmune disease programs. (**a**) Analytical strategy for intersecting ChromBPNet-predicted functional variants with LCL *cis*-eQTLs. (**b**) Upset plot displaying the intersection among different levels of variant functionality. caQTL variants and MPRA emVars were derived from LCLs (see Extended Data Fig. 7b-d). (**c**) Rank-ordered plot of 332 genes associated with 783 ChromBPNet-predicted functional fine-mapped LCL eQTL variants. (**d**) Distribution of 783 ChromBPNet-predicted functional eQTL variants across chromosomes. Gray shaded regions indicate the proportion for chr6 and chr17 variants that map to the HLA or 17q21.31 locus, respectively. (**e**) Bar plot displays distribution of 481 ChromBPNet-predicted functional eQTL variants across autoimmune diseases. Variants residing in the HLA or 17q21.31 locus were omitted from the analysis.

